# Develop a durable, memory-driven, CspZ-targeting Lyme disease vaccine by rationale adjuvant selection

**DOI:** 10.64898/2026.01.08.698480

**Authors:** Miranda McCarty, Sergio A. Hernández, Jill Malfetano, Ana Carolina Leão, Maria Jose Villar, Xiuli Yang, Yi-Lin Chen, Jungsoon Lee, Zhuyun Liu, Ulrich Strych, Maria Elena Bottazzi, Utpal Pal, Klemen Strle, Wen-Hsiang Chen, Yi-Pin Lin

## Abstract

Rational adjuvant selection is a systematic approach based on adjuvant-mediated immunomodulation to identify safe vaccine regimens that enhance protective immunity. Transmitted through ticks and caused by the bacterium *Borrelia burgdorferi* (*Bb*), Lyme disease (LD) is the most common vector-borne disease in the Northern hemisphere. There are no effective vaccines, making it suitable for testing the concept of rational adjuvant selection. Here, we formulated our previously developed and effective LD vaccine antigen, CspZ-YA_C187S,_ with different adjuvants suitable for human use; we analyzed the immune response by transcriptomics and tested the vaccine efficacy after *Bb* infection. We identified Alum-CpG and Alum-αGal to elicit the highest titers of CspZ-YA_C187S_-dependent protective antibodies and robust levels of protection but through distinct mechanisms of immunomodulation. We demonstrated that immunization with Alum-CpG formulated CspZ-YA_C187S_ provided up to nine months of protective bactericidal antibody titers, as well as recall-memory response to prevent LD after natural infection. Immunity was linked to elevated levels of IgG1 memory cells in the vaccine-triggered immune responses. This work thus identified a durable, memory immunity-driven LD vaccine, ultimately paving the road to understanding the mechanisms of rationale adjuvant selection for vaccine development.

## INTRODUCTION

Vaccines depend on the host’s adaptive immune response to confer protection. An ideal vaccine offers safe, robust, and long-lasting protective immune responses. Each component of the vaccine, namely the antigen and the adjuvant, triggers distinct immune responses, but not all responses persist and/or confer protection ^1, 2^. Some immune responses may even promote pathogen dissemination, exacerbate disease, and/or pose safety concerns ^3^. Consequently, considerable efforts have focused on optimizing vaccine regimens to prolong protective immune responses while minimizing adverse effects ^4^. One practical approach of vaccine regimen optimization is rational adjuvant selection whereby antigens and adjuvants are selected and combined based on their capacity to shape and elicit a favorable immune response ^5, 6^. Broadly, adjuvants can act as either delivery systems or immune potentiators, according to their mechanism of action ^7^. Delivery systems (e.g., oil-water emulsion, microparticles, or minerals such as aluminum hydroxide (Alum)) modulate antigen release to enhance durability of protection and promote antigen-specific responses ^8^. Immune potentiators activate specific responses due to engagement of receptors and agonism of particular pathways (e.g., α-Galactosylceramide (αGal) for invariant natural killer T (iNKT) cells; PAM3CSK4 (PAM3) for Toll-like receptors (TLR) 1 and 2; CpG oligodeoxynucleotides (CpG) for TLR-9) ^7^. Knowing which particular immune responses are required for pathogen survival or disease prevention allows for adjuvants to be selected individually or combinatorially to direct host immune responses toward pathogen clearance and mitigation of pathology. Understanding how specific adjuvant mechanisms translate into protective immunity is, therefore, central to rational adjuvant selection.

Lyme disease is the most common vector-borne infection in the Northern Hemisphere, with more than 476,000 human cases reported in the United States annually ^9^. Transmitted by *Ixodes* ticks, this disease is caused by a variety of bacterial species within the *Borrelia burgdorferi* sensu lato complex (also known as *Borreliella burgdorferi* or Lyme borreliae) ^1^. LYMErix, the only licensed human Lyme disease vaccine to date, was withdrawn from the market 20 years ago ^10, 11^. This vaccine targeted a bacterial protein present in ticks only, OspA, or outer surface protein A, to block transmission to humans ^10, 11^. A next-generation OspA-based vaccine, VLA15, is currently in clinical trials ^12, 13, 14^. However, because Lyme borreliae express *ospA* predominantly in the tick midgut and downregulate *ospA* expression after transmission to mammals, to sustain high titers of anti-OspA antibodies required for protection, typically, higher vaccine doses and multiple boosters are necessary ^11, 13, 15, 16, 17^. Efforts were taken to attempt in addressing such caveats through optimizing adjuvants and antigens in regimens^18, 19, 20, 21, 22, 23^. Although the tick-specific expression of *ospA* likely does not induce OspA-specific memory responses after natural infection in humans, the history of OspA vaccine development suggests Lyme disease as a tractable system for elucidating the immunological basis of how adjuvant formulations optimize protective and durable immunity.

The strategy we have pursued to overcome the challenges of Lyme disease vaccine development is to identify new vaccine antigens. Lyme borreliae produce outer surface proteins that bind host Regulators of Complement Activation (RCAs) to promote the degradation of the complement protein, C3b ^24, 25, 26, 27^. This ultimately facilitates complement resistance and survival of the bacteria in circulation despite high concentrations of complement. Five of these Lyme borreliae proteins, so-called Complement Regulator Acquiring Surface Proteins (CRASPs), bind to a host RCA, Factor H (FH) ^24^. Among them, CspZ (aka CRASP-2) is essential for optimal bacterial dissemination after invasion of vertebrate hosts ^20, 28^, consistent with the fact that CspZ is produced only when bacteria are in mammalian hosts, but not in ticks ^29^. Furthermore, anti-CspZ IgG is constantly detectable in serologically confirmed Lyme disease patients and infected murine models ^30, 31, 32, 33^, supporting CspZ as an antigen target for Lyme disease vaccine development. However, because vaccination with wild-type CspZ has shown limited protection in mice ^20, 31, 34, 35^, we have engineered a superior mutated antigen, CspZ-YA_C187S_ (CspZ-Y207A/Y211A/C187S)^20, 33, 35^. By exposing potential bactericidal epitopes near FH-binding sites via Y207A/Y211A mutations and stabilizing overall folding via the C187S mutation, CspZ-YA_C187S_ has been shown to effectively prevent *B. burgdorferi* colonization and joint inflammation with fewer immunization boosters^20, 33, 35^. These efforts enable us to apply rational adjuvant selection by further identifying CspZ-YA_C187S_-induced host immune responses and linking these responses to durable and memory-mediated protection against Lyme disease.

In this study, we formulated CspZ-YA_C187S_ with different adjuvants established for clinical use and identified (i) the resulting immunomodulation of each formulation, (ii) the efficacy of vaccination in a murine challenge model for Lyme borreliae, and (iii) the durability and memory-inducing characteristics of the protective antibodies triggered by these formulations following natural infection.

## RESULTS

### CpG or αGal boost IgG titers and bactericidal activity triggered by Alum-CspZ-YA_C187S_ vaccination

Amongst the adjuvant used in humans HogenEsch, ^36, 37^, Alum(Aluminum hydroxide/Alhydrogel) can trigger a Th2-biased immune response and antibody production and IL-10, which can alliviate Lyme disease-associated arthritis preclinically ^38^. PAM3(PAM3CSK4) facilitates TLR2-mediated cytokines documented to promote *B. burgdorferi*-specific recognition by the host for clearance ^39^. To define vaccine formulations that elicit the highest levels of CspZ-YA_C187S-_specific IgG titers and protective antibodies, we thus immunized mice once or twice on days 0 and 14 with CspZ-YA_C187S_ formulated with Alum and PAM3. IgG titers after 14 and 28 days post initial immunization (dpii) were compared to those generated by formulating CspZ-YA_C187S_ with TiterMax Gold (TMG), an adjuvant previously used in preclinical mouse studies ^20, 33, 35^ (**Fig. 1A**). Anti-CspZ-YA_C187S_ IgG titers were similar across all groups after one or two immunizations (**Fig. 1B-C**). We then quantified bactericidal activity against *B. burgdorferi* strain B31-A3 (OspC type A, the most common genotype from patients in North America) using BA_50_ values, the serum dilution that kills 50% of spirochetes (**Fig. 1D-G, Table S1**). After a single immunization, killing was detectable for the TMG formulation, but not for the Alum or PAM3 vaccines (**Fig. 1D-E, Table S1**). Following the second immunization, TMG adjuvanted CspZ-YA_C187S_ produced significantly higher BA_50_ values than Alum or PAM3 formulations (**Fig. 1F-G, Table S1**), indicating that Alum or PAM3 alone do not outperform the TMG benchmark.

**Figure 1.**
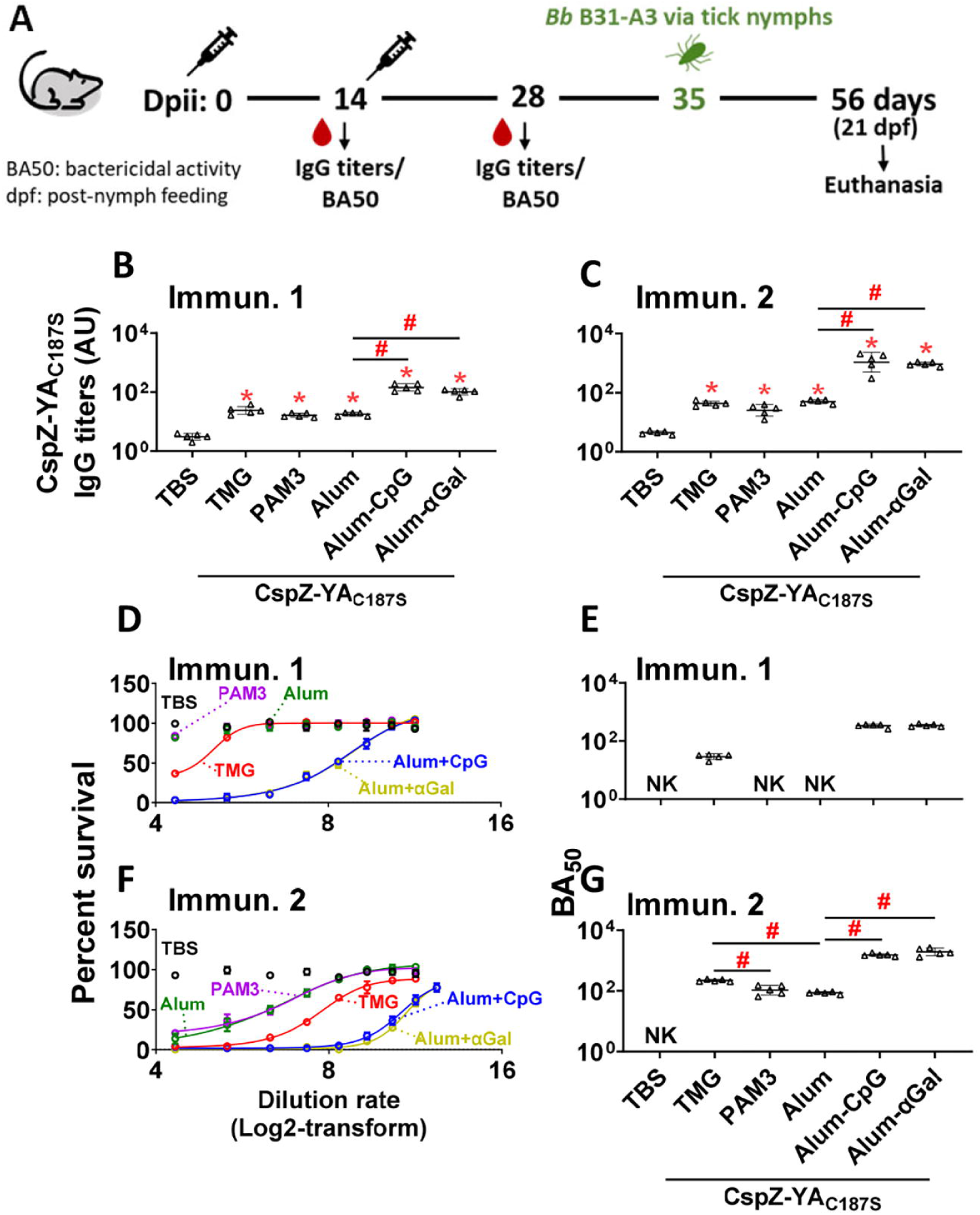
Vaccination with CspZ-YA_C187S_ formulated with Alum-CpG or Alum-αGal induced robust antibody titers and bactericidal activities in mice. **(A)** Pre-adolescent C3H/HeN mice received two inoculations at 0- and 14-days post initial immunization (dpii) with TBS (control) or CspZ-YA_C187S_ formulated with the indicated adjuvants, followed by infection via feeding of *I. scapularis* nymphs carrying *B. burgdorferi* strain B31-A3 at 35 dpii and euthanasia at 56 dpii. Sera were collected at **(B, D, E)** 14 dpii for one immunization (Immun. 1) and **(C, F, G)** 28 dpii for two immunizations (Immun. 2). **(B-C)** The levels of total IgG against CspZ-YA_C187S_ in the sera were determined using quantitative ELISA. Data shown are the geometric mean ± geometric standard deviation of the titers of anti-CspZ-YA_C187S_ antibodies from five mice per group. **(D-G)** Sera were diluted as indicated, and mixed with guinea pig complement and *B. burgdorferi* B31-A3 for 24 hours. Surviving spirochetes were quantified microscopically from three fields of view in three independent experiments. **(D, F)** Survival percentage was derived from the proportion of serum-treated to untreated spirochetes. Data shown are the mean ± SEM of the survival percentage from three replicates in one representative experiment. (E, G) The BA_50_ value, representing the dilution rate that effectively killed 50% of spirochetes, was obtained from curve-fitting and extrapolation of Panels C and F. Data shown are the geometric mean ± geometric standard deviation of the borreliacidal titers from five mice per group in three experiments per mouse sample and shown in Table S1. (“NK”), no killing. Statistical significance (p < 0.05, Kruskal Wallis test with the two-stage step-up method of Benjamini, Krieger, and Yekutieli) of IgG titers or borreliacidal titers from an indicated group of mice was compared to the titers from TBS-inoculated mice (“*”) or between indicated groups were shown as “#”.

Alum primarily functions as a delivery system for antigen formulations and has been commonly co-formulated with other adjuvants considered “immune potentiators” to enhance pathogen-killing efficacy ^7^. We thus next evaluated whether adding immune potentiators could enhance responses. As a one of the immune potentiators, αGal facilitates iNKT cells/B cells production, the immune mechanisms documented to promote Lyme borreliae clearance and prevent Lyme disease-associated joint inflammation preclinically ^40, 41^. The other immune potentiator, CpG, is a agonist of TLR9, reported to be critical for Lyme borreliae clearance ^42, 43^. We thus co-formulated Alum-CspZ-YA_C187S_ with CpG or αGal (**Fig. 1A**). Both combinations of immune potentiators significantly increased anti-CspZ-YA_C187S_ IgG titers relative to Alum alone after one or two immunization doses (**Fig. 1B-C**). Importantly, bacterial killing was detectable after a single immunization with Alum-CpG or Alum-αGal formulated CspZ-YA_C187S_ and indistinguishable from those with TMG formulated CspZ-YA_C187S_ (**Fig. 1D-E, Table S1**) and were significantly higher than Alum alone after the booster (**Fig. 1F-G, Table S1**). Together, these data demonstrate that adding CpG or αGal as immune potentiators to Alum augments both antibody titers and *B. burgdorferi*-killing.

### CpG or αGal formulations with Alum-CspZ-YA_C187S_ prevent *B. burgdorferi* infection after two immunizations

After the mice were immunized with CspZ-YA_C187S_ formulated with various adjuvants, they were challenged by feeding of *B. burgdorferi* B31-A3-infected *Ixodes scapularis* nymphal ticks at 35 dpii (**Fig. 1A**). Uninfected mice or TBS-inoculated and *B. burgdorferi*-infected mice were included as controls. Bacterial burdens in replete nymphs were comparable across all vaccine groups, agreeing with prior reports that mouse vaccination with TMG-CspZ-YA_C187S_ does not impact spirochetal burdens in fed nymphs (**Fig. 2A**) ^33^. Seropositivity to the *B. burgdorferi* C6 peptide at 21 days post-nymph feeding (dpf) mirrored protection: all TMG-CspZ-YA_C187S_ animals remained C6-seronegative, whereas all mice in the TBS control and Alum-formulated groups, and most of the PAM3-formulated group (5/5 mice for TBS and Alum; 4/5 for PAM3), were C6-seropositive. Mice immunized with TBS, Alum, or PAM3 formulations had significantly higher anti-C6 IgG titers than uninfected controls **(Fig. 2B)**. In contrast, all mice receiving either Alum-CpG or Alum-αGal formulation were seronegative to the C6 peptide, similar to uninfected animals **(Fig. 2B)**. Consistent with serology, bacterial burdens at the tick bite site, bladder, heart, and knee joints were elevated in the TBS, Alum-, and PAM3-formulated groups but were undetectable in uninfected controls, TMG, Alum-CpG, and Alum-αGal groups (**Fig. 2C-F**). Thus, CpG and αGal adjuvants convert Alum-CspZ-YA_C187S_ into an efficacious vaccine that blocks *B. burgdorferi* tissue colonization and seroconversion preclinically.

**Figure 2.**
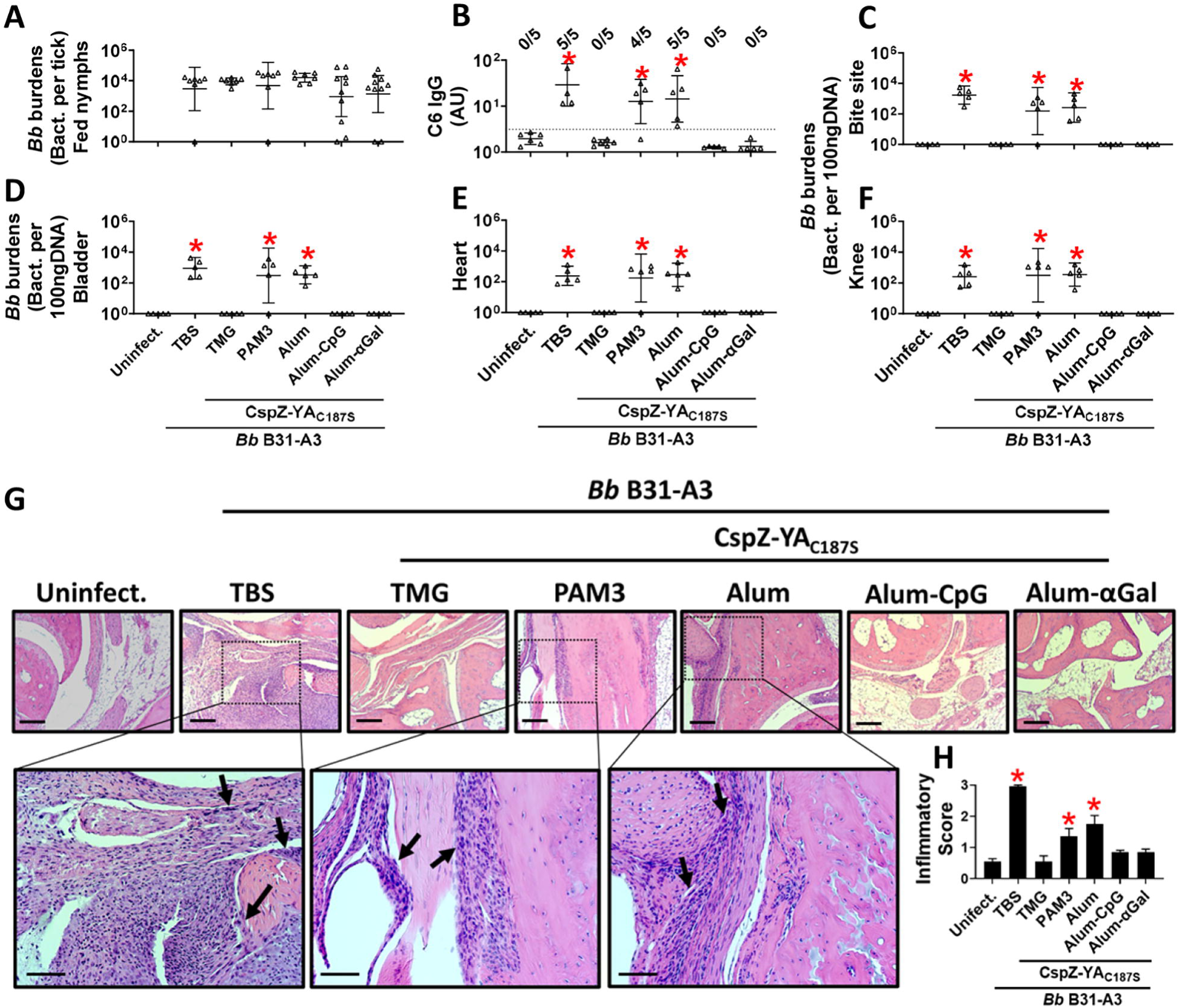
CspZ-YA_C187S_ formulated with Alum-CpG or Alum-αGal protected mice from seroconversion, borrelial colonization and Lyme disease-associated joint inflammation. Five pre-adolescent C3H/HeN mice were inoculated with TBS or CspZ-YA_C187S_ formulated with indicated adjuvants, followed by infection using **(A)** nymphs carrying *B. burgdorferi* B31-A3 in the fashion described in Fig. 1A. Mice inoculated with TBS that are not fed on by nymphs were included as an uninfected control group (uninfect.). **(B)** Seropositivity was determined by measuring the levels of IgG against C6 peptides in the sera of those mice at 21 dpf using ELISA. A mouse was considered as seropositive if that mouse had IgG levels against C6 peptides greater than the threshold, the mean plus 1.5-fold standard deviation of the IgG levels against C6 peptides from the TBS-inoculated, uninfected mice (dotted line). The number of mice in each group with anti-C6 IgG levels greater than the threshold (seropositive) is shown. Data presented here are the geometric mean ± geometric standard deviation of the titers of anti-C6 IgG. Statistical significances (p < 0.05, Kruskal-Wallis test with the two-stage step-up method of Benjamini, Krieger, and Yekutieli) of differences in IgG titers relative to (*) uninfected mice are presented. (A, C-F) *B. burgdorferi* (*Bb*) burdens at **(A)** nymphs after feeding to repletion or **(C**) the tick feeding site (“Bite Site”), **(D)** bladder, **(E)** heart, and **(F)** knees, were quantitatively measured at 21 dpf, shown as the number of (A) *Bb* per tick or **(C-F)** per 100 ng total DNA. Data shown are the geometric mean ± geometric standard deviation of the spirochete burdens from each group of mice. Asterisks indicate the statistical significance (p < 0.05, Kruskal Wallis test with the two-stage step-up method of Benjamini, Krieger, and Yekutieli) of differences in bacterial burdens relative to uninfected mice. **(G-H)** Tibiotarsus joints at 42 dpli were collected to assess inflammation by staining these tissues using hematoxylin and eosin. **(G)** Representative images from one mouse per group are shown. Top panels are lower-resolution images (joint, ×10 [bar, 160 µm]); bottom panels are higher-resolution images (joint, 2×20 [bar, 80 µm]) of selected areas (highlighted in top panels). Arrows indicate infiltration of immune cells. **(H)** To quantitate inflammation of joint tissues, at least ten random sections of tibiotarsus joints from each mouse were scored on a scale of 0-3 for the levels of inflammation. Data shown are the mean inflammation score ± standard deviation of the inflammatory scores from each group of mice. Asterisks indicate the statistical significance (p < 0.05, Kruskal Wallis test with the two-stage step-up method of Benjamini, Krieger, and Yekutieli) of differences in inflammation relative to uninfected mice.

We next determined the severity of *B. burgdorferi*-triggered inflammation in the tibiotarsal (ankle) joints of mice at 21 dpf (**Fig. 1A**). Histopathology revealed dense mononuclear infiltrates in the muscles, tendons, and connective tissues from all five mice immunized with TBS or Alum-, and most (4/5) PAM3-immunizeed mice (arrows in representative images, **Fig. 2G**). Inflammation scores in these vaccination groups were significantly elevated versus uninfected controls (**Fig. 2H**). Conversely, all five mice immunized with CspZ-YA_C187S_ formulated with Alum-CpG or Alum-αGal displayed no pathologic joint inflammation and had inflammatory scores comparable to uninfected controls and the protective TMG adjuvant formulation (**Fig. 2G-H**). Overall, these data show that the addition of CpG or αGal to Alum-CspZ-YA_C187S_ vaccination yields robust protection after two immunizations, preventing seroconversion, tissue colonization, and Lyme disease-associated joint inflammation, whereas PAM3 or Alum does not.

### Alum-CpG and Alum-αGal formulations drive distinct immune transcriptional programs after CspZ-YA_C187S_-vaccination

We next compared the transcriptomic programs induced by adding CpG or αGal to Alum-formulated CspZ-YA_C187S._ RNA was isolated from whole spleens of mice 14 days after the second immunization (28 dpii) and profiled for RNA sequencing (RNA-seq) (**Fig. 1A**). Relative to TBS controls, mice vaccinated with Alum-CpG-formulated CspZ-YA_C187S_ produced fewer differentially expressed genes (DEGs) whereas Alum-αGal-formulated CspZ-YA_C187S_ yielded more DEGs overall, suggesting distinct gene expression profiles induced by CpG or αGal in Alum-CspZ-YA_C187S_ immunized mice (**Fig. S1, Dataset 1**). Using Alum-CspZ-YA_C187S_ mice as the baseline, Alum-CpG vs. Alum identified 8 upregulated DEGs, whereas Alum-αGal versus Alum identified 100 upregulated DEGs. Only three DEGs (2.9%) were shared between these upregulated sets (**Fig. 3A**). In contrast, for downregulated DEGs, Alum-CpG and Alum-αGal yielded 28 and 32 downregulated genes, respectively, with no overlap (**Fig. 3B**). In the direct transcriptomic comparison of the Alum-CpG vs. Alum groups, we observed the upregulation of genes involved mostly in immunoglobulin production and selected inflammatory mediators (**Fig. 3C-D, Table S2**). In Alum-αGal vs. Alum, we observed upregulation of immunoglobulin genes together with antigen presentation/recognition, Th-1 cell modulation, and leukocyte trafficking, accompanied by reduced expression of several anti-inflammatory genes (**Fig. 3E-F, Table S3**). Taken together, these results suggest that CpG and αGal remodel splenic transcriptomics in qualitatively different ways when formulated with Alum-CspZ-YA_C187S_.

**Figure 3.**
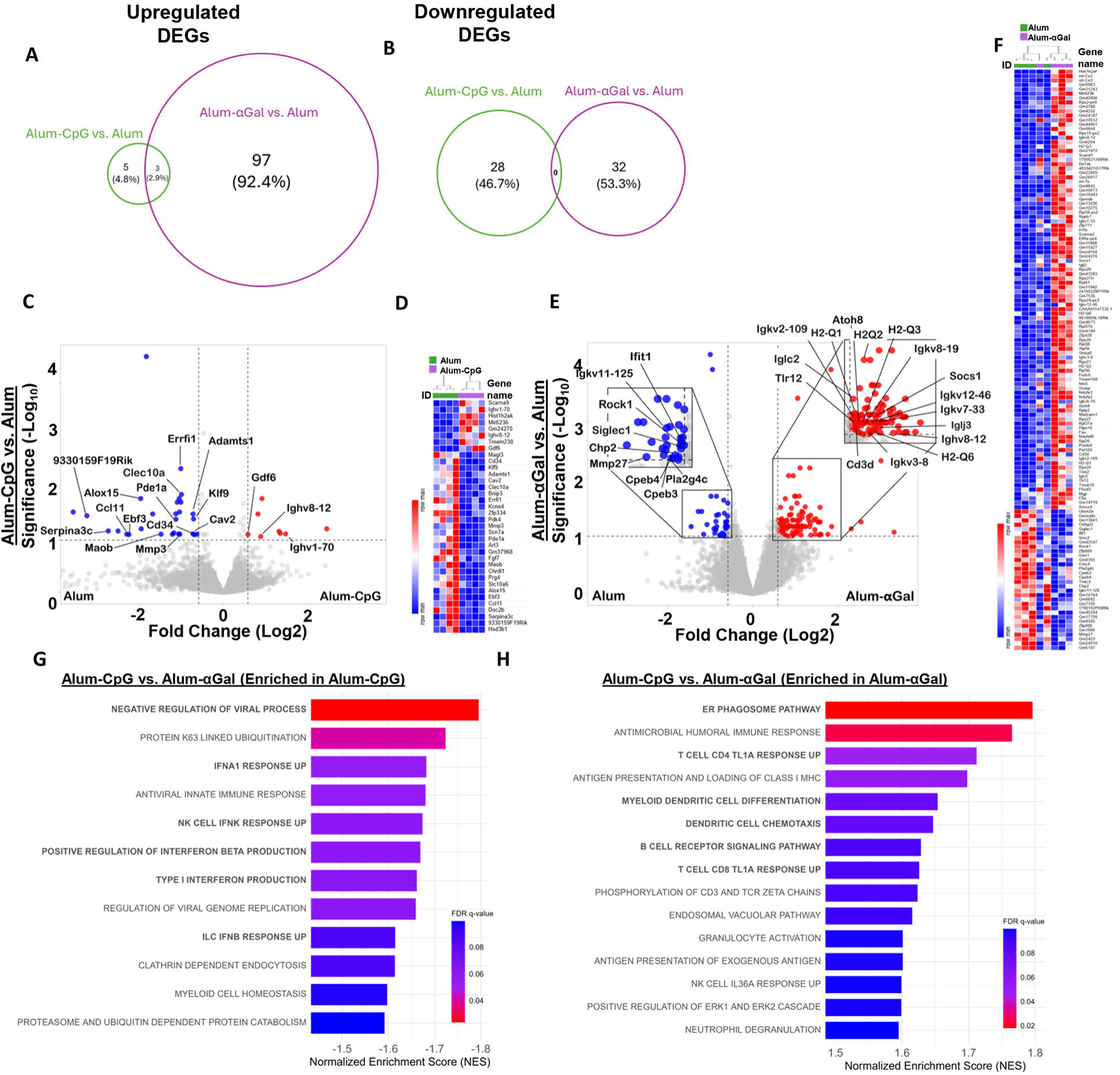
CpG or αGal skewed the Alum-CspZ-YA_C187S_-induced immune pathways in different fashions. Five pre-adolescent C3H/HeN mice per group were inoculated with TBS or CspZ-YA_C187S_ formulated with Alum, Alum-CpG, and Alum-αGal twice in the fashion described in Fig. 1A**. (A-H)** Spleens were collected at 28dpii, and the expression levels of genes in the spleen from each group of the mice were determined. Differentially expressed genes (DEGs) are defined by an adjusted P values less than 0.05 and absolute (log_2_fold change) ≥ 0.58 or ≤ -0.58. **(A-B)** Venn diagram comparing the number of **(A)** up- or **(B)** down-regulated DEGs identified in the spleen from mice inoculated with CspZ-YA_C187S_ formulated with Alum-CpG vs. Alum and Alum-αGal vs. Alum. **(C, E)** The expression levels of genes in the comparison of **(C)** Alum-CpG vs. Alum and **(E)** Alum-αGal vs. Alum are plotted as a Volcano plots. The up- and down-regulated DEGs are shown in red and blue, respectively. The immune-related DEGs in each comparison are highlighted. The details of all DEGs are shown in **Dataset S1**, and the details of these immune related DEGs are shown in **Table S2 to S3**. **(D, F)** Heat map exhibits the expression patterns of all DEGs in the comparison of **(D)** Alum-CpG vs. Alum and **(F)** Alum- αGal vs. Alum, and the gene IDs are shown in the right. **(G-H)** GSEA analysis to identify the enriched pathways triggered by the CspZ-YA_C187S_ formulated with Alum-CpG or Alum-αGal when the enrichment analysis was performed in the comparison of the formulations under Alum-CpG vs. Alum-αGal. The DEGs enriched in the formulation of **(G)** Alum-CpG vs. **(H)** Alum- αGal are shown. The pathways shown were based on the statistical cutoff (FDR q-value <0.1 and NES >1.5).

We next conducted pathway/over-representation analyses modulated by CpG or αGal when added to Alum-CpG-CspZ-YA_C187S_-vaccination. Given the modest DEG lists, pathway/term over-representation tests were underpowered. We therefore utilized Gene Set Enrichment Analysis (GSEA) on the complete ranked gene lists, enabling detection of modest but relevant pathway changes with statistical control (False Discovery Rate (FDR)). In the comparisons vs. Alum, immune-related functions were prominent among the top 10 significantly enriched pathways in both formulations (3 of the top 10 in Alum-CpG; 4/10 in Alum-αGal) (bold pathways in **Fig. S2**). We thus selected all immune-related pathways significantly upregulated in the mice vaccinated with CspZ-YA_C187S_ formulated with Alum-CpG or Alum-αGal (vs. Alum-only) and characterized their functions. Most of the highest-ranked immune terms in both groups (6/14 in Alum-CpG; 8/19 in Alum-αGal) mapped to adaptive B and T cell functions (**Fig. S3**), aligning with the significantly greater levels of IgG titers and enhanced bactericidal activity seen for the Alum-CpG and Alum-αGal formulations of CspZ-YA_C187S_ (**Fig. 1B-G**). To further differentiate the immune transcriptional programs induced by the formulations containing Alum-CpG and Alum-αGal, we directly compared the pathways upregulated by the two adjuvants relative to each other using GSEA. Alum-CpG (enriched relative to Alum-αGal) favored many interferon (IFN)-related pathways, including IFN-α responses, positive regulation of IFN-β production, NK-cell IFN-κ signatures, and an antiviral state overall (see bold pathways in **Fig. 3G**). In contrast, Alum-αGal (enriched relative to Alum-CpG) emphasized antigen processing and presentation, as well as antigen presenting cells (APC)/lymphocyte crosstalk pathways (see bold pathways in **Fig. 3H**). In sum, CpG skews Alum-CspZ-YA_C187S_ towards an antiviral/interferon-modulated program, whereas Alum-αGal induces APC licensing, antigen processing, and lymphocyte priming processes. These results suggest two distinct routes of immunity for the two formulations that lead to a robust, protective humoral responses after CspZ-YA_C187S_ vaccination.

### Alum-CpG vs. Alum-αGal impact cytokine secretion in *ex vivo* splenocytes from CspZ-YA_C187S_ immunized mice but converge on memory B-cell activation

We isolated splenocytes from mice at 7-days after the second inoculation with TBS (control), or CspZ-YA_C187S_ formulated with Alum, Alum-αGal or Alum-CpG. Splenocytes were restimulated *ex vivo* with CspZ-YA_C187S_, and cytokine supernatant concentrations were quantified. Across all measured mediators (IL-10, TNFα, IL-2, IL-6, IL-4, IFN-γ at two doses), splenocytes from Alum-αGal-vaccinated mice produced significantly higher levels than TBS controls (**Fig. 4A-G**). In contrast, vaccination with Alum-CpG adjuvant selectively increased only IFN-γ levels, while other cytokines were not significantly different (**Fig. 4F and G**), supporting our transcriptomic findings of Alum-CpG amplifying a marked IFN-specific immune program, whereas Alum-αGal-CspZ-YA_C187S_ elicits a broader cytokine profile.

**Figure 4.**
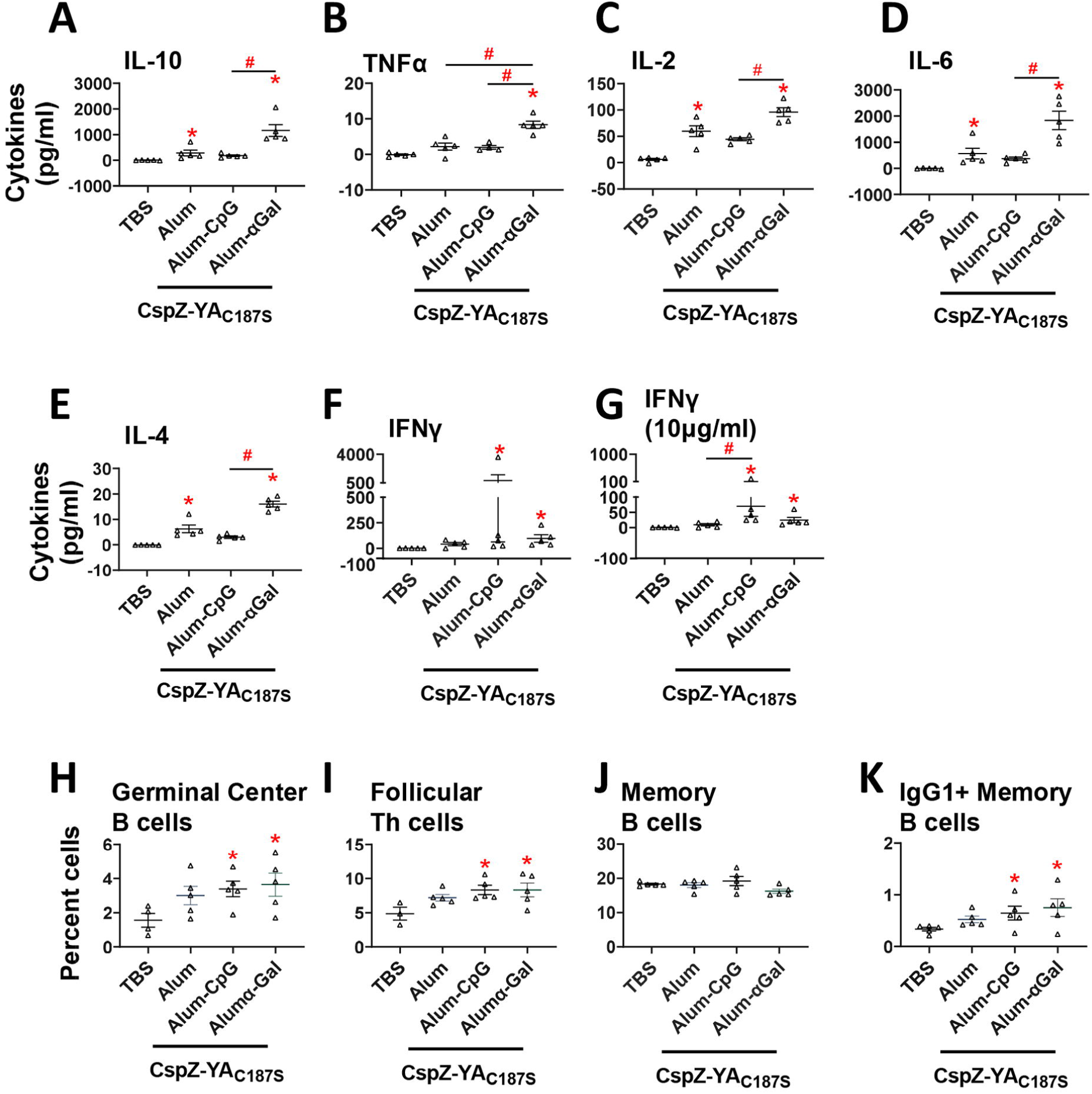
The splenocytes from vaccination with Alum-CpG or Alum-αGal formulated CspZ-YA_C187S_ differed in the elevated cytokine species but were similar in the activated memory cell types. Splenocytes isolated from the mouse spleens at 21 dpii were restimulated with CspZ-YA_C187S_ for 48-h. **(A-G)** Multiplex Luminex analysis was used to measure the indicated secreted cytokines. **(H-K)** Flow cytometry analysis was applied to determine the percentage of **(H)** Germinal Center B cells (CD19+CD38-FAS+), **(I)** Follicular T helper cells (CD4^+^CD44^+^CXCR5hiPD^-^1hi), **(J)** Memory B cells (CD19^+^CD38^+^IgD^-^IgM^-^), and **(K)** IgG1+ Memory B cells (CD19^+^CD38^+^IgD^-^IgM^-^IgG1^+^). Data shown are the mean ± SEM of the **(A-G)** cytokine levels or **(H-K)** percentage of live/singlets cell populations from each group of mice. The statistical significance (p < 0.05, Kruskal Wallis test with the two-stage step-up method of Benjamini, Krieger, and Yekutieli) of differences in bacterial burdens relative to TBS-inoculated mice (“*”) or between indicated groups of mice (“#”) were indicated.

To delineate the responders among splenocytes to the CspZ-YA_C187S_ vaccination with different tested adjuvants, we profiled splenocytes by flow cytometry after antigen (CspZ-YA_C187S_) restimulation. Germinal center (GC) B cells and T follicular helper (Tfh) cells were significantly increased in mice immunized with Alum-CpG or Alum-αGal but not in mice immunized with Alum alone, compared to TBS-inoculated mice (**Fig. 4H-I**). Total memory B cell populations were similar among the tested groups of splenocytes, but IgG1+ memory B cells were significantly enriched in both Alum-CpG and Alum-αGal cohorts compared to the TBS-inoculated mice (**Fig. 4K**). These findings indicate that although CpG and αGal diverge in the acute cytokine profile they elicit, both adjuvants promote GC B cells/Tfh engagement, and expansion of IgG1+ memory B cells, consistent with the enhanced antibody titers and protective phenotypes reported earlier.

### Vaccination with Alum-CpG-CspZ-YA_C187S_ elicits durable, bactericidal antibodies for up to nine months

The abovementioned results identified Alum-CpG and Alum-αGal as the lead adjuvants to formulate with CspZ-YA_C187S_. However, CpG, but not αGal, has been FDA-approved and used in licensed human vaccines ^44, 45, 46, 47^, we thus prioritized the Alum-CpG formulation and sought to determine the durability of protective antibodies triggered by CspZ-YA_C187S_ vaccination. Mice received either one dose (at day 0) or two doses (at day 0 and 14dpii). Serum was collected longitudinally at different days post last immunization (dpli) to quantify antigen-specific IgG titers against OspA and CspZ-YA_C187S_, as well as bactericidal activity (BA_50_) (**Fig. 5A**). TBS-inoculated mice and mice immunized three times with TMG-OspA were included as controls (**Fig. 5A**). We detected anti-OspA IgG only in the TMG-OspA group, with antibodies persisting until 182 dpli. No OspA reactivity was observed in any mice inoculated with Alum-CpG-CspZ-YA_C187S_ or TBS (**Fig. 5B and S4**). Conversely, TMG-OspA or TBS-inoculated mice did not exhibit detectable anti-CspZ-YA_C187S_ IgG, but these specific antibodies were present in high titers in Alum-CpG-CspZ-YA_C187S_ (**Fig. 5C and S5**). A two-dose immunization regimen with CspZ-YA_C187S_ yielded signifianctly higher titers at all time points and extended seropositivity to 238 dpli, compared with 210 dpli after a single dose (**Fig. 5C and S5**). No bactericidal activity was detected in TBS-inoculated mice. The TMG-OspA immunized mice maintained measurable bactericidal antibody titers (BA_50_) until 182 dpli (**Fig. 5D and S6**). Importantly, sera from Alum-CpG-CspZ-YA_C187S_ immunized mice showed durable bactericidal activity up until 210 dpli after one dose and up to 238 dpli after two doses, mirroring the IgG durability (**Fig. 5D and S6**). These results showed the capacity of Alum-CpG-CspZ-YA_C187S_ to induce dose-responsive, long-lived humoral immunity with bactericidal function lasting up to nine months, supporting this formulation as the lead vaccine formulation.

**Figure 5.**
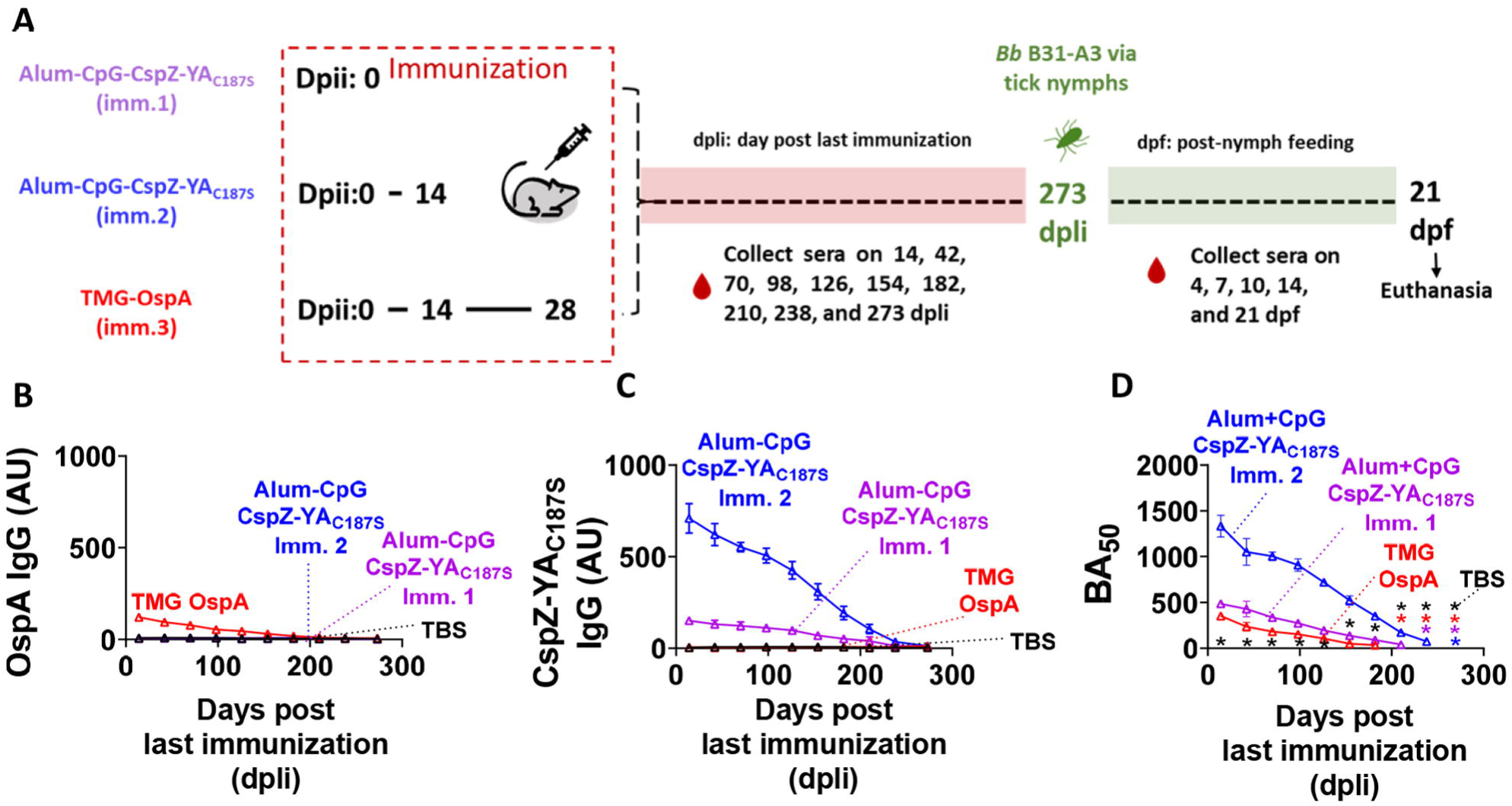
Titers and bactericidal activities of antibodies triggered by Alum-CpG formulated CspZ-YA_C187S_ were detectable for up to nine months. **(A)** The schematic diagram shows pre-adolescent C3H/HeN mice receiving one or two injections of CspZ-YA_C187S_ formulated with Alum-CpG on 0 and 14 dpii.. Mice inoculated with TBS at 0 and 14 dpii, or TMG-formulated OspA at 0, 14, and 28dpii were included as control. Sera were collected at 14, 42, 70, 98, 126, 154, 182, 210, 238, and 273 days post last immunization (dpli). **(B-C)** The levels of total IgG against **(B)** OspA and **(C)** CspZ-YA_C187S_ in the sera were determined using quantitative ELISA. Data shown are the geometric mean ± geometric standard deviation of the titers of **(B)** anti-OspA and **(C)** CspZ-YA_C187S_ IgGs from five mice per group. **(D)** The sera were diluted as indicated and mixed with guinea pig complement and *B. burgdorferi* B31-A3 for 24 hours. Surviving spirochetes were quantified from three fields of view microscopically in three independent experiments. The survival percentage was derived from the proportion of serum-treated to untreated spirochetes. Data shown are the mean ± SEM of the survival percentage from three replicates in one representative experiment. The BA_50_ value, representing the dilution rate that effectively killed 50% of spirochetes, was obtained from curve-fitting and shown in **Table S4**. Asterisks represent that no killing was detected in indicated groups of mice and time points.

### Natural infection rapidly recalled bactericidal anti-CspZ antibodies and conferred protection in Alum-CpG-CspZ-YA_C187S_-vaccinated mice

We next determined whether natural infection could recall CspZ-targeting immunity once vaccine-induced CspZ-YA_C187S_ IgG waned below detection. Mice previously immunized with Alum-CpG-CspZ-YA_C187S_ (one or two doses), TMG-OspA (three doses), or TBS were held until 273 dpli, when anti-CspZ-YA_C187S_ IgG and bactericidal antibody titers (BA_50_) were undetectable (**Fig. 5A**). Animals were then exposed to *B. burgdorferi* by infected nymphal tick feed and sera were collected at 4, 7, 10, 14, and 21 dpf to assess antibody recall (**Fig. 5A**). As expected, anti-OspA IgG remained negligible across all immunization groups after tick feeding, including mice who received TMG-OspA (**Fig. 6A, S7**), consistent with *ospA* downregulation in vertebrate hosts ^15, 16^. In contrast, Alum-CpG-CspZ-YA_C187S_ vaccination primed a rapid recall response, as anti-CspZ-YA_C187S_ IgG rose significantly compared to uninfected controls by 4-7 dpf and remained elevated through all timepoints (**Fig. 6B, S8**). TMG-OspA and TBS groups lacked early anti**-**CspZ-YA_C187S_ increases, and showed only modest responses by 10 dpf, in line with detectable CspZ-targeting antibodies reported at later stages during natural infection (*de novo* priming) ^20, 29, 30, 32, 33^(**Fig. 6B, S8**). Antibody isotyping at 7 dpf, done to determine early protective responders, revealed a broad recall of IgG subtypes (IgG1, IgG2a, IgG2b, IgG3) (**Fig. 6C**). Among these isotypes, IgG1 was found to be the predominant subclass (**Fig. 6C**), mirroring the increase in IgG1+ memory B cells observed post-vaccination (**Fig. 4K**). Further, early bactericidal function was unique to sera from Alum-CpG-CspZ-YA_C187S_ recipients, as BA_50_ became detectable at 4-7 dpf and increased thereafter, whereas other groups lacked bactericidal activity until ≥10 dpf (**Fig. 6D, S9, Table S4**). These results suggest an early emergence of memory immunity-driven, bactericidal antibodies after natural infection in the mice immunized with Alum-CpG-formulated CspZ-YA_C187S._

**Figure 6.**
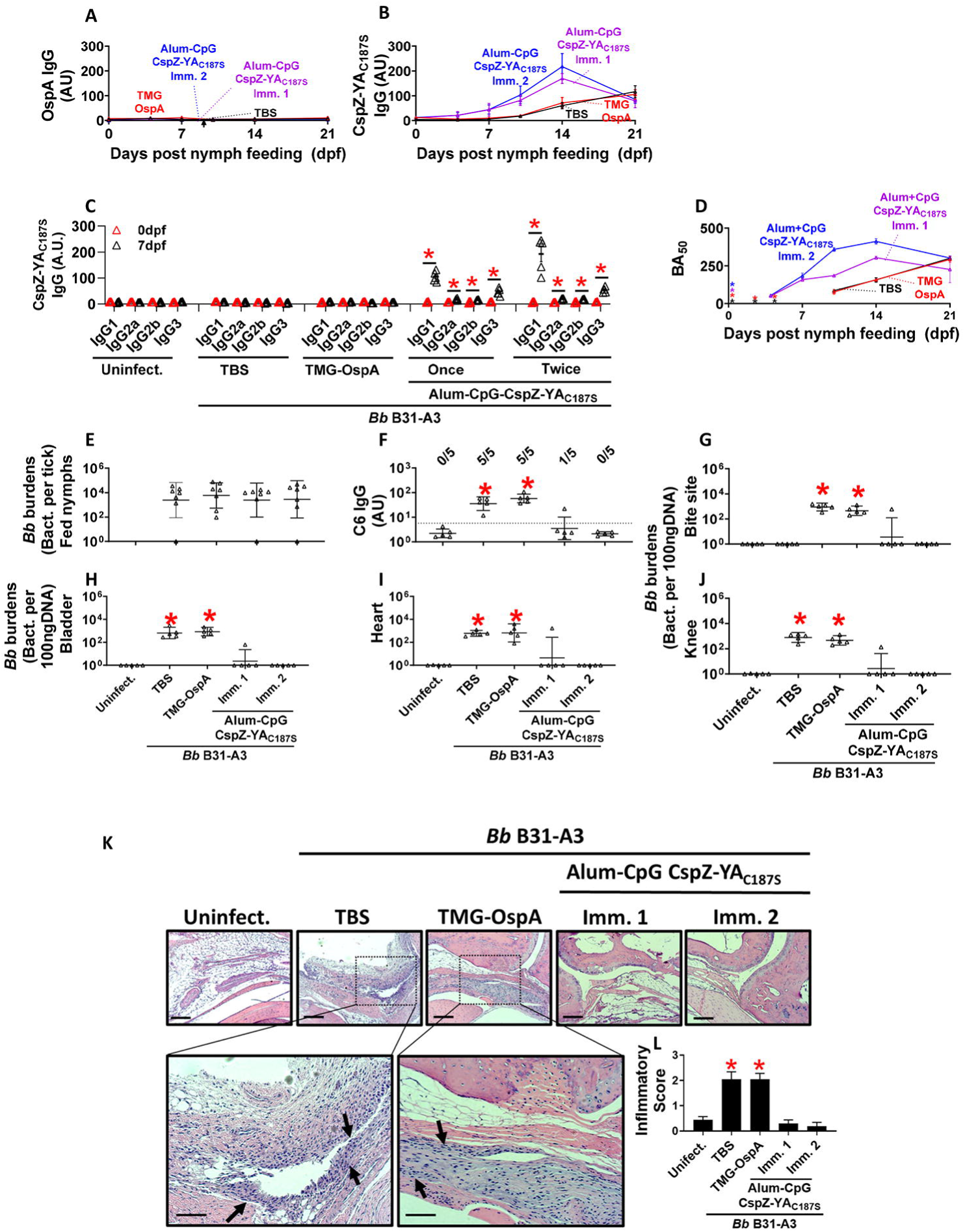
Natural infection triggered CspZ-targeting bactericidal antibodies linked to prevention of seroconversion, borrelial colonization and joint inflammation in Alum-CpG-CspZ-YA_C187S_ vaccinated mice. As shown in Fig. 5A, five pre-adolescent C3H/HeN mice were inoculated with TBS or CspZ-YA_C187S_ formulated with Alum-CpG. At 273 dpli, using *I. scapularis* nymphs carrying *B. burgdorferi* strain B31-A3, the mice were naturally infected. Sera were collected at 4, 7, 10, 14, and 21 dpf, and mice were euthanized at 21 dpf. Mice inoculated with TBS that were not fed on by nymphs were included as an uninfected control group (uninfect.). **(A-C)** The levels of total IgG against **(A)** OspA and **(B)** CspZ-YA_C187S_ or indicated IgG isotypes against **(C)** CspZ-YA_C187S_ in the sera at indicated time points were determined using quantitative ELISA. Data shown are the geometric mean ± geometric standard deviation of the titers of **(A)** anti-OspA and **(B-C)** CspZ-YA_C187S_ IgGs from five mice per group. **(D)** The sera were diluted as indicated, and mixed with guinea pig complement and *B. burgdorferi* B31-A3 for 24 hours. Surviving spirochetes were quantified from three fields of view microscopically in three independent experiments. The survival percentage was derived from the proportion of serum-treated to untreated spirochetes. Data shown are the mean ± SEM of the survival percentage from three replicates in one representative experiment. The BA_50_ value, representing the dilution rate that effectively killed 50% of spirochetes, was obtained from curve-fitting and shown in **Table S5**. Asterisks represent that no killing was detected in indicated groups of mice and time points. **(E)** The *B. burgdorferi burdens* of nymphs carrying *B. burgdorferi* B31-A3. Mice inoculated with TBS that were not fed on by nymphs were included as an uninfected control group (uninfect.). **(F)** Seropositivity was determined by measuring the levels of IgG against C6 peptides in the sera of those mice at 21 dpf using ELISA. The mouse was considered as seropositive if that mouse had IgG levels against C6 peptides greater than the threshold, the mean plus 1.5-fold standard deviation of the IgG levels against C6 peptides from the TBS-inoculated, uninfected mice (dotted line). The number of mice in each group with the anti-C6 IgG levels greater than the threshold (seropositive) is shown. Data shown are the geometric mean ± geometric standard deviation of the titers of anti-C6 IgG. (G-J) *B. burgdorferi* (*Bb*) burdens at **(G**) the tick feeding site (“Bite Site”), **(H)** bladder, **(I)** heart, and **(J)** knees, were quantitatively measured at 21 dpf, shown as the number of (E) *Bb* per tick or **(G-J)** per 100ng total DNA. Data shown are the geometric mean ± geometric standard deviation of the spirochete burdens from each group of mice. **(K-L)** Tibiotarsus joints at 21 dpf were collected to assess inflammation by staining these tissues using hematoxylin and eosin. **(K)** Representative images from one mouse per group are shown. Top panels are lower-resolution images (joint, ×10 [bar, 160 µm]); bottom panels are higher-resolution images (joint, 2×20 [bar, 80 µm]) of selected areas (highlighted in top panels). Arrows indicate infiltration of immune cells. **(L)** To quantitate inflammation of joint tissues, at least ten random sections of tibiotarsus joints from each mouse were scored on a scale of 0-3 for the levels of inflammation. Data shown are the mean inflammation score ± standard deviation of the inflammatory scores from each group of mice. Asterisks indicate the statistical significance (p < 0.05, Kruskal Wallis test with the two-stage step-up method of Benjamini, Krieger, and Yekutieli) of differences in **(C, E, G-J)** bacterial burdens, **(F)** C6 IgG titers, **(L)** inflammatory scores relative to uninfected mice.

We next tested if this rapid recall via CspZ-YA_C187S_ vaccination translated into protection by determining anti-C6 seropositivity, *B. burgdorferi* burdens in tissues, and ankle joint inflammation histologically at 21 dpf (**Fig. 5A**). Tick bacterial burdens in fed nymphs did not differ among groups, as expected (**Fig. 6E**), but host outcomes differed sharply. At 21 dpf, all TBS- or TMG-OspA inoculated mice were C6-seropositive and harbored significantly higher *B. burgdorferi* burdens in skin (bite site), bladder, heart, and knee, compared to uninfected mice (**Fig. 6F-J**). These mice also displayed extensive mononuclear infiltrates in ankle joints and significantly elevated histologic inflammation scores relative to uninfected controls (**Fig. 6K-L**). In contrast, one mouse out of five immunized with a single dose of Alum-CpG-CspZ-YA_C187S_ and none in the two-dose group became C6-seropositive, with bacterial tissue burdens indistinguishable from those of uninfected mice (**Fig. 6F-J**). Similarly, one animal (single dose) and none from the two-dose group developed apparent joint inflammation, resulting in histopathological scores comparable to those from uninfected mice (**Fig. 6J-K**). Taken together, these results link a durable, memory B-cell antibody response primed by Alum-CpG formulated CspZ-YA_C187S_ vaccination and rapidly recalled by natural infection, to the prevention of C6-seropositivity, bacterial colonization, and *B. burgdorferi*-associated joint pathology in the murine Lyme disease model.

## DISCUSSION

Adjuvants are critical in vaccines to enhance magnitude, quality, and durability of the protective immunity against infectious diseases. For many vaccine prototypes, rational adjuvant selections that utilize established immune potentiators or delivery systems to trigger desirable immune pathways while avoiding unwanted host responses can improve immunization safety and efficacy ^5, 6^. Guided by this principle, we evaluated how different immune potentiators layered onto a common delivery system (Alum) direct diverse immune responses elicited by a structure-guided CspZ vaccine (CspZ-YA_C187S_). Lyme borreliae are positioned as a tractable model pathogen for vaccine development, given the numerous efforts invested to elucidate host responses that enable pathogen clearance and alleviate the development of manifestations ^25^. These data provide the foundation for applying the concept of rational adjuvant selection to elicit disease-resolving immune responses in the development of Lyme disease vaccines. As the most commonly used adjuvant in human vaccines, Alum activates the local innate immune system through pattern recognition receptors (PRRs), which matures antigen presenting cells (APCs) for appropriate presentation of the vaccine antigen. This process layered to immune potentiators ultimately creates a unique inflammatory niche that produces CspZ-YA_C187S_ antibodies to drive more robust bactericidal responses, while reducing the number of boosters required ^48, 49, 50^. However, CspZ-YA_C187S_ formulated with Alum alone did not provide protection in our study. Additionally, it is well-known that TLR1/2 recognize Lyme borreliae, triggering inflammatory pathways favoring spirochete elimination ^51, 52, 53, 54, 55^ and that PAM3 is a TLR1/2 agonist ^56, 57^. Yet, similar to Alum, PAM3-formulated CspZ-YA_C187S_ also failed to protect mice from *B. burgdorferi* colonization or joint inflammation after immunization. These findings likely reflect insufficient priming of protective humoral and cellular immune programs by Alum or PAM3 adjuvants alone. We thus tested immune potentiators with defined mechanisms co-formulated with Alum to bias an immune response towards protective pathways that drive beneficial adaptive immune responses. One of the selected immune potentiators is CpG, a TLR9 agonist known to trigger APC signaling, drives activation of immune cells and promotes Th1 immune responses, and proliferation and maturation of B cells ^58, 59, 60, 61^. Another immune potentiator, α-galactosylceramide (αGal), is a synthetic glycolipid that can be presented by CD1d, which is widely expressed on hematopoietic and non-hematopoietic cells, and subsequently recognized by iNKT cells ^62, 63^. Consistent with TLR9 ligation and iNKT cells in facilitating Lyme borreliae clearance ^40, 41, 42, 64, 65, 66^, both adjuvant combinations (Alum-CpG and Alum-αGal) protected against *B. burgdorferi* infection in murine models following vaccination with CspZ-YA_C187S_. Additionally, mice immunized with each of these two formulations developed significantly higher levels of anti-CspZ-YA_C187S_ IgG and borreliacidal activities than animals vaccinated with Alum-CspZ-YA_C187S_ alone, supporting well-documented engagement of downstream B cell responses by TLR9 and iNKT cell pathways ^67, 68, 69, 70^. Our results thus suggest similar favorable immune responses (i.e., higher extents of bactericidal antibodies) could be boosted by distinct pathways using diverse immune potentiators, underscoring the need to identify the unique mechanisms of potentiator-mediated protective immunity.

In this study, transcriptomic profiling of splenocytes provided additional mechanistic insights into specific adaptive immune responses of the immune simulators. Compared to TBS-inoculated mice, both Alum-CpG and Alum-αGal-formulated CspZ-YA_C187S_ upregulated genes associated with B cell and T cell immunity, including the activation of signaling, immunoglobulin production, and effector functions with associated cytokines. This is in line with previous findings of the requirement of durable T-dependent B cell immune responses for effective Lyme borreliae clearance ^49, 71^. Furthermore, the enhanced expression of both T and B cell associated genes can be directly attributed to CpG and αGal priming, as these genes remained significantly upregulated relative to mice immunized with CspZ-YA_C187S_ formulated with Alum alone. However, Alum-αGal formulated CspZ-YA_C187S_ uniquely permitted the enrichment of NK cell-related pathways, agreeing with αGal as an iNKT cell agonist ^60, 61, 62, 63^. Alum-αGal also preferentially enriched antigen-processing and presentation programs (e.g., ER phagosome pathway and MHC assembly/loading) in APCs (dendritic cells), together with the aforementioned iNKT/NK-cell-linked terms. These results suggest the possibility for the role of αGal in promoting more efficient APC licensing and crosstalk with T cells, thereby broadening adaptive immune responses for Lyme borreliae clearance. Further, we found that immunization with Alum-CpG formulated CspZ-YA_C187S_ demonstrated an interferon-centric signature. Activation of IFN-specific pathways was corroborated with targeted proteomics using splenocytes from mice immunized with Alum-CpG formulated CspZ-YA_C187S_. These consistent results agree with published data on TLR9-triggered type IFN (IFN-α and IFN-β) signaling ^72^. Together with type II IFNs (i.e., IFN-γ) and IFN-γ-induced cytokines being associated with lower *B. burgdorferi* burdens in humans and the phenotypes of *B. burgdorferi* clearance in the murine models ^66, 73, 74, 75^, our findings provide possible mechanisms for Alum-CpG-activated IFN immunity in protection against Lyme borreliae colonization. Despite these upstream differences between the formulations of Alum-CpG and Alum-αGal, both regimens converged on expansion of germinal center B cells, T follicular helper cells, and IgG1+ memory B cells, all aligning with increased antibody titers and serum bactericidal activity. Moreover, TLRs, including TLR9, signal through MyD88, which can result in a broad spectrum of cytokine secretion ^76, 77^. However, we did not observe a lot of pathways other than those IFN-associated be significantly enriched by Alum-CpG formulated CspZ-YA_C187S_ when compared to Alum-αGal-CspZ-YA_C187S_, suggesting the possibility of antigen (CspZ-YA_C187S_)-adjuvant interplay in immunomodulation. This is consistent with many Lyme borreliae lipoproteins reported as having immunomodulatory functions ^34, 51, 78, 79^. Overall, the findings here provide mechanistic information on how adjuvants can skew host protective immune responses.

The induction of immune memory is critical for vaccine development as the capacity and durability of these responses impact the frequency of immunization needed to be effective ^80^. For Lyme disease vaccines, the ideal response would include the capability to trigger long-lasting memory after immunization and rapid emergence of bactericidal antibodies upon natural infection ^81^. However, the currently developed OspA-based vaccine blocks the transmission of the *B. burgdorferi* to the host but lack the ability to trigger persisting OspA-targeted antibodies after several months (unless specific formulations were used ^10, 11, 23^). Additionally, in support of the downregulation of OspA after *B. burgdorferi* invades vertebrate hosts^20, 28^, we found no induction of OspA-specific IgG in OspA-immunized mice after natural infection. Unlike OspA, Alum CpG–CspZ YA_C187S_ generated CspZ specific IgG and borreliacidal antibodies for up to nine months in murine models, with higher magnitude and longer persistence after two doses than one. This persistence of CspZ IgG exceeds what is typically achieved with OspA-based vaccine, suggesting that CspZ-targeted vaccines may provide longer lasting immunity and reduce the numbers of required booster doses. We also found that CspZ-targeted responses were readily recalled, with Alum-CpG-primed animals mounting a rapid memory response detectable as early as 4-days after natural infection. Such rapid responses were accompanied by a rise in functional bactericidal activity, protection from tissue colonization, seroconversion, and joint inflammation. Antibody isotype and flow cytometry analyses point to a mechanistic link between vaccine-primed memory B cell responses and protection ^82^. The dominance of IgG1 at recall and the expansion of IgG1+ memory B cells after immunization with Alum-CpG formulated CspZ-YA_C187S_ support a model in which early reactivation of class-switched memory, likely aided by Alum-CpG-driven interferon signaling and efficient T-cell help, delivers complement-fixing antibodies. This posits IgG1 as the major contributor of CspZ-YA_C187S_-primed memory immunity which confers protection. IgG isotype differences against *B. burgdorferi* antigens have been linked to infection vs. protection after *B. burgdorferi* invasion ^49, 83^. For some antigens (i.e., OspC), IgG1 has been associated with protection, consistent with this subclass preferentially triggering classical complement pathway-mediated pathogen killing, which has been documented in Lyme borreliae clearance ^84, 85^. Thus, our work supports the role of IgG1 in Lyme disease protection, not only induced by vaccines but also in memory immune responses following natural infection. Overall, IgG1-mediated protective responses from CspZ-YA_C187S,_ or other previously identified Lyme borreliae vaccine antigens, could be the determinant of protection upon natural infection. Further examination of this possibility has the potential to aid development of more precise antibody-based protection in memory immune responses induced by Lyme disease vaccines.

We also showed that unvaccinated mice or mice immunized with other regimens that do not confer protection to natural infection (i.e., TMG-OspA) also develop anti-CspZ-YA_C187S_ IgG. This result is consistent with the elevated titers of CspZ-specific IgG in Lyme disease patients and mice colonized by Lyme borreliae ^30, 31, 32, 33^. However, mice vaccinated with CspZ-YA_C187S_ had an earlier onset of CspZ-specific IgG antibodies, compared to the unvaccinated or TMG-OspA-immunized mice (4 dpf vs. 10 dpf). This result highlights the importance of a rapid recall of anti-CspZ-YA_C187S_ IgG for an efficacious CspZ-targeted vaccine. In fact, Lyme borreliae infection kinetics begin with the pathogens transiently being in the bloodstream, followed by fast migration to distal tissues ^86, 87^. Such a unique progression during infection is largely due to the vulnerability of Lyme borreliae while in the bloodstream, in which numerous host mechanisms are present to eradicate spirochetes ^25, 88^. Although Lyme borreliae develop several antibody-evasion strategies immediately, those strategies do not mature until later in the infection ^89, 90, 91^. Therefore, the ability to generate bactericidal antibodies within 4 days during natural infection, at a time when spirochetes are transiently blood-borne and most susceptible to humoral clearance, is expected to be critical in CspZ-YA_C187S_ vaccine efficiency. Taken together, our study tested the concept of various adjuvant formulations in eliciting beneficial memory B cell immune responsesto CspZ-YA_C187S_. The optimization of adjuvant selection identified a formulation that can facilitate a long-lasting, memory-driven Lyme disease vaccine. Overall, our work examines the concept of rational adjuvant selection for vaccine development and elucidates the vaccine-triggered, adjuvant-influenced immune responses, which would benefit the development of other infectious disease vaccines.

## MATERIALS AND METHODS

### Ethics Statement

All mouse experiments were performed in strict accordance with all provisions of the Animal Welfare Act, the Guide for the Care and Use of Laboratory Animals, and the PHS Policy on Humane Care and Use of Laboratory Animals. The protocols, docket numbers 22-451 and G2024-07, were approved by the Institutional Animal Care and Use Agency of Wadsworth Center, New York State Department of Health and Tufts University, respectively. All efforts were made to minimize animal suffering.

### Mouse, ticks, and bacterial strains

Four-week-old, female C3H/HeN mice were purchased from Charles River (Wilmington, MA, USA). Although such an age of the mice has not reached sexual maturity, the under development of immune system in this age of mice would allow such mice to be more susceptible to Lyme borreliae infection, increasing the signal to noise ratio of the readout. That will also provide more stringent criteria to define the protectivity. BALB/c C3-deficient mice were from in-house breeding colonies ^92^ and *Ixodes scapularis* tick larvae were obtained from BEI Resources (Manassas, VA). *Escherichia coli* strain BL21(DE3) and derivatives were grown at 37°C or other appropriate temperatures in Luria-Bertani broth or agar, supplemented with kanamycin (50µg/mL). *B. burgdorferi* strain B31-A3 were grown at 33°C in BSK II complete medium ^93^. Cultures of *B. burgdorferi* B31-A3 were tested with PCR to ensure a full plasmid profile ^94, 95^.

### Purification of OspA and CspZ-YA_C187S_

The purification of hexahistidine-tagged lipidated OspA and untagged CspZ-YA_C187S_ was executed as described ^33, 96, 97^. The *E. coli* BL21(DE3) strains producing OspA and CspZ-YA_C187S_ are listed in **Table S6**. OspA was lipidaded as lipidation is required for recombinant OspA proteins to protect mice from Lyme disease infection^97, 98^.

### Mouse immunization and infection

C3H/HeN mice were immunized as described, with slight modifications ^33^. Twenty-five µg of untagged CspZ-YA_C187S_ or hexahistine-tagged lipidated OspA (control) in 50 µl of Tris-buffered saline (TBS) buffer was thoroughly mixed with 50 µl TiterMax Gold adjuvant (Norcross, GA, USA), resulting 100µl inoculum. Twenty-five µg of tagged CspZ-YA_C187S_ was also formulated with PAM3 (10 μg, PAM3CSK4, InvivoGen, CA, USA), Alum (400 μg, Alhydrogel, Corda, NJ, USA), Alum (400 μg) mixed with CpG (20 μg, CpG1826, InvivoGen, CA, USA), or Alum (400 μg) mixed with αGal (1 μg, α-galactosylceramide, Avanti, AL, USA) in 50 µl of **TBS** with 0.05% Pluronic F-127 and 7% sucrose (TBS). In addition, 50 µl of TBS only was included as a control. The inoculum was introduced into C3H/HeN mice subcutaneously once at 0- or twice at 0- and 14-days post initial immunization (**Fig. 1A and 4A**). Blood was collected via submandibular bleeding at 14-days post last immunization (dpli) (**Fig. 1A**) or 14, 42, 70, 98, 126, 154, 182, 210, 238, and 273 dpli (**Fig. 5A**). Sera obtained from the blood were used to determine the IgG titers of CspZ-YA_C187S_ or OspA, and the bactericidal activities, described in the section “ELISAs” and “Bactericidal assays”, respectively. At 21 dpli (35 days post initial immunization shown in **Fig. 1A**) or 273 dpli (**Fig. 5A**), *B. burgdorferi* B31-A3-infected flat nymphs were placed in a chamber on C3H/HeN mice as described ^34^. Five nymphs were allowed to feed to repletion on each mouse, and a subset of nymphs was collected pre- and post-feeding. At 21 days post nymph feeding (dpf), tick bite sites of skin, bladder, knees, and heart were collected to determine the bacterial burdens, and ankles were also collected at 21 dpli to determine the severity of arthritis described in the section “Quantification of spirochete burdens and histological analysis of arthritis.” At this time point, blood was also collected via cardiac puncture bleeding to isolate sera for the determination of seropositivity described in the section “ELISAs.”

### ELISAs

The determination of the titers of anti-OspA, CspZ-YA_C187S_ IgG, C6 IgG or the IgG isotypes (IgG1, IgG2a, IgG2b, and IgG3) in the serum samples was as described, with modifications ^20, 33, 81^. Specifically for anti-C6 IgG, this methodology has been commonly used for human Lyme disease diagnosis ^99^. One µg of untagged and lipidated OspA, untagged CspZ-YA_C187S_, or C6 peptides was coated on ELISA plate wells as described ^20^. The procedures following the protein coating are as described previously ^20^. For each serum sample, the maximum slope of optical density/minute of all the dilutions of the serum samples was multiplied by the respective dilution factor, and the greatest value was used as arbitrary unit (AU) to represent the antibody titers for the experiment to obtain anti-OspA, -CspZ-YA_C187S_, or -C6 IgG, or IgG1, IgG2a, IgG2b, and IgG3 against CspZ-YA_C187S_. For anti-C6 IgG, the seropositive mice were defined as the mice with the serum samples yielding a value greater than the threshold, the mean plus 1.5-fold standard deviation of the IgG values derived from the uninfected mice.

### Borreliacidal assays

The ability of serum samples to eradicate *B. burgdorferi* B31-A3 was determined as described with modifications ^20, 33, 35^. Briefly, the sera collected from mice immunized with lipidated OspA or CspZ-YA_C187S_ under different adjuvant formulations at different immunization frequency were heat-treated to inactivate complement. The rest procedure was performed in the similar fashion as described ^20, 35^. Surviving spirochetes at 24 h of incubation with Guinea pig sera (Sigma) and different dilution rate of serum samples were quantified by directly counting the motile spirochetes using dark-field microscopy and expressed as the proportion of serum-treated to untreated Lyme borreliae. The 50% borreliacidal activities (BA_50_), representing the serum dilution rate that kills 50% of spirochetes, was calculated using dose-response stimulation fitting in GraphPad Prism 9.3.1.

### Quantification of spirochete burdens and histological analysis of arthritis

DNA was extracted from the indicated mouse tissues to determine the bacterial burdens using quantitative PCR analysis as described ^20, 33^. Basically, the forward and reverse primers with the sequences as GTGGATCTATTGTATTAGATGAGGCTCTCG and GCCAAAGTTCTGCAACATTAACACCTAAAG, respectively, were used to amplify the *recA* gene of *B. burgdorferi* strain B31-A3. The number of *recA* copies was calculated by establishing a threshold cycle (Cq) standard curve of a known number of *recA* gene extracted from strain B31-A3, and burdens were normalized to 100 ng of total DNA from mouse tissues or presented as the number of bacteria per tick for *B. burgdorferi* in ticks. The histological analysis was applied to ankles for Lyme disease-associated arthritis (**Fig. 3G**), as described ^20, 33^. Basically, images were scored based on the severity of the inflammation as 0 (no inflammation), 1 (mild inflammation with less than two small foci of infiltration), 2 (moderate inflammation with two or more foci of infiltration), or 3 (severe inflammation with focal and diffuse infiltration covering a large area).

### mRNA Extractions for RNASeq analyses

Spleens were placed immediately on dry ice and stored at –80 C° and then submitted for RNA extraction and RNAseq analysis by GENEWIZ (Waltham, MA). Turbo Capture mRNA Kits (Qiagen) were used for total RNA extraction of 50 mg of tissues. A drill and polypropylene pestles were used to lyse TRIzol® suspended tissues in Eppendorf Tubes, followed by the treatment of DNAse I as described in the vendor’s manuals. Elutions were prepared in DNAase and RNAase free water, and the quality of RNA was measured using the Nanodrop Eight Spectrophotometer (ThermoFisher Scientific), targeting a 260/280 ratio of 1.95-2.04. RINe (RNA Integrity Number) values greater than 6.5 with strong indications of robust 18S and 28S (28S/18S) peaks, with minimal evidence of degraded small RNA at the lower threshold range, were submitted for further processing. The RNA samples were processed for mRNA enrichment using Dynabead’s mRNA Purification Kit per vendor’s manual (ThermoFisher). The mRNA sequencing was then carried out by polyA selection using Illumina HiSeq, PE 2x150 (150 bp paired end).

### RNASeq analysis

After the sequencing, the reads were trimmed via Trimmomatic v.0.36, and mapped to the *Mus musculus* strain C3H/HeJ reference genome via ENSEMBL using the STAR aligner v.2.5.2b. The gene hit counts (calculation of reads/gene/sample) were determined using feature counts from the Subread package v.1.5.2. The results from Salmon were imported into R v.4.2.3 with tximport and a DESeq2 object was created with the DESeq dataset from tximport function. Differential expression analyses were performed by DESeq2 with minreplicates for replace set to 30. The identification of DEGs was shown in **Dataset S1**. RNASeq results from the spleens of mice immunized with CspZ-YA_C187S_ formulated with Alum, Alum-CpG, or Alum-αGal were first compared with those from the control (TBS-inoculated mice). Additionally, the results from the spleens of mice vaccinated with Alum-CpG or Alum-αGal were compared with those from mice injected with Alum. VolcaNoseR was used to plot and explore differentially expressed genes (DEGs) in volcano plot format ^100^. Adjusted p-values ≤ 0.05 and log2 fold changes ≥ 0.58 or ≤ -0.58 were used as the initial cutoffs defining differentially expressed genes (DEG). The number shared and unique DEGs among the mice immunized with CspZ-YA_C187S_ formulated Alum, Alum-CpG, or Alum-αGal was visualized as Venn diagram with Venny 2.1 ^101^. Heat map analysis was performed using Morpheus ^102^. The Pearson correlation was used to measure dissimilarities between individuals for the heat map’s hierarchical clustering ^102^. Complete ranked gene lists were analyzed with GSEA (v4.4.0; http://software.broadinstitute.org/gsea/index.jsp), using 1,000 gene-set permutations, minimum/maximum gene-set sizes of 15/500, respectively ^103^. We queried the following mouse MSigDB collections (v2025.1): M2 (curated pathways), M5 (ontology), M7 (immunologic signatures), and M8 (cell-type signatures). Gene sets were considered enriched at FDR q <0.25 with normalized enrichment scores (NES) |NES| ≥ 1.5. The NES sign indicates direction of enrichment (positive = comparator group, negative = baseline).

### Multiplex analysis of cytokines by Luminex

Splenocytes from mice immunized with CspZ-YA_C187S_ formulated with each of the tested adjuvants were prepared and analyzed by Luminex, as described previously ^104^. Spleens were rinsed and homogenized in sterile PBS buffer, using a gentleMACS Dissociator (Miltenyi Biotech, Charlestown, MA) and treated with ACK lysis buffer (Lonza, Portsmouth, NH) to remove red blood cells. The suspension was then diluted 5-fold with complete Roswell Park Memorial Institute (RPMI) 1640 medium (cRPMI, Sigma Aldrich) supplemented with 10% fetal bovine serum (FBS, Sigma Aldrich), 1× penicillin-streptomycin (Pen-Strep, Sigma Aldrich), and L-glutamine (Sigma Aldrich), followed by centrifugation at 300 × g for 5 min. Splenocytes were resuspended in 5 mL of cRPMI, filtered through 40 µm strainers (BD Biosciences, San Jose, CA) and counted using acridine orange– propidium iodide (AOPI, Fisher Scientific) staining on a Cellometer Auto 2000 (Nexcelom Bioscience, Lawrence, MA). For stimulation assays, viable splenocytes (1 × 10^6^) were incubated for each sample in a 96-well non-tissue culture plate and re-stimulated with different concentrations of CspZ-YA_C187S_ or cRPMI (control). Splenocytes were incubated at 37°C in 5% CO_2,_ and cell-free supernatants were collected after 48-h after re-stimulation. Cytokines levels of IL-10, TNF-a, IL-2, IL-6, IL-4, and IFN-g in supernatants were measured by a Luminex-based assay using Milliplex MAP Mouse Th 17 Magnetic Bead Panel (Millipore Sigma), as previously described ^105^. Cytokine responses were calculated by subtracting values obtained from splenocytes restimulated with cRPMI alone from those restimulated with CspZ-YA_C187S_.

### Flow Cytometry of splenocytes

Splenocytes from the section “Multiplex analysis of cytokines by Luminex” were washed with 1X PBS, and stained with Live/Dead NIR fixable viability dye (ThermoFisher). The viable population of the cells were incubated with purified rat anti-mouse CD16/CD32 (Mouse BD Fc Block) and stained by the antibodies indicated below obtained from ThermoFisher. Basically, the population of Tfh cells was stained with and evaluated using Alexa Fluor 700 conjugated anti-CD4 IgG, fluorescein isothiocyanate (FITC) conjugated anti-CD44 IgG, BV421 Brilliant Violet 421 (BV-421) conjugated anti-CD185 (for CXCR5) IgG and PE (phycoerythrin)-Cy7 (PE-Cy7) conjugated anti-CD279 (PD-1) IgG. To evaluate the population of B cells, splenocytes were stained with FITC conjugated anti-CD4, anti-CD8 FITC, anti-Ly-6G/Ly-6C (Gr-1), anti-F4/ 80, or anti-CD11c IgG, BV-421 or BV-711 conjugated anti-CD19 IgG, Cy5.5 or BV-711 conjugated anti-CD38 peridinin chlorophyll protein (PerCP) IgG, BV-605-conjugated anti-mouse CD95 (Fas) IgG, allophycocyanin (APC) conjugated anti-CD138 IgG, Alexa Fluor 700 conjugated anti-IgD, and/or PE-Cy7 conjugated anti-IgM, and/or PerCP-Cy5.5 conjugated anti-IgG1. Stained cell samples were then applied to an Attune Flowcytometer (ThermoFisher), and at least 100,000 total events in a live gate were analyzed using FlowJo software. Cells were gated on forward and side scatter for lymphocytes, exclusion of viability dye, and singlet populations. B cells and Tfh cells in the spleen were determined based on Fluorescence minus one (FMO).

### Statistical analyses

Significant differences were determined with a Kruskal-Wallis test with the two-stage step-up method of Benjamini, Krieger, and Yekutieli ^106^ and two-tailed Fisher test (for seropositivity in **Fig. 2B and 5B**) ^107^ using GraphPad Prism 9.3.1. A p-value < 0.05 was used to determine significance.

## AUTHOR CONTRIBUTIONS

M.C.: Conceptualization, Methodology, Validation, Formal analysis, Investigation, Data Curation, Writing - Original Draft, Writing - Review & Editing, Visualization. S.H.: Conceptualization, Software, Methodology, Validation, Formal analysis, Investigation, Data Curation, Writing - Review & Editing, Visualization. J.M.: Methodology, Investigation. A.C.L.: Methodology, Validation, Formal analysis, Investigation, Data Curation, Writing - Review & Editing, M.J.V.: Methodology, Validation, Formal analysis, Investigation, Data Curation, Writing - Review & Editing, X.Y.: Methodology, Investigation, Data Curation, Y.L.C.: Investigation, Writing - Review & Editing, J.L.: Methodology, Investigation, Writing – Review & Editing, Z.L.: Investigation, Writing - Review & Editing, U.S.: Methodology, Investigation, Writing - Review & Editing, M.E.B.: Conceptualization, Methodology, Investigation, Writing - Review & Editing, Supervision, Project administration, Funding acquisition. U.P.: Supervision, Project administration, K.S.: Methodology, Investigation, Supervision, Project administration, Funding acquisition, W.H.C.: Conceptualization, Methodology, Validation, Formal analysis, Investigation, Data Curation, Writing - Original Draft, Writing - Review & Editing, Visualization, Supervision, Project administration, Funding acquisition, Y.P.L.: Conceptualization, Methodology, Validation, Formal analysis, Investigation, Data Curation, Writing - Original Draft, Writing - Review & Editing, Visualization, Supervision, Project administration, Funding acquisition.

## ACKNOWLEDGEMENTS

The authors thank Patricia Rosa and John Leong for providing *B. burgdorferi* strain B31-A3. We thank Carly Fernandes and Connor McKaig’s technical support. We are also grateful for Kate Sulka in helping to review the manuscript and provide advice. We thank the Wadsworth Animal Core for assistance with Animal Care and Tufts Laboratory Animal Medicine Service.

## DATA AVAILABILITY

All data generated or analysed during this study are included in this published article and its supplementary information files.

