## Supplementary Information for "Develop a durable, memory-driven, CspZ-targeting Lyme disease vaccine by rationale adjuvant selection"

**SUPPLEMENTAL FIGURES AND FIGURE LEGENDS**

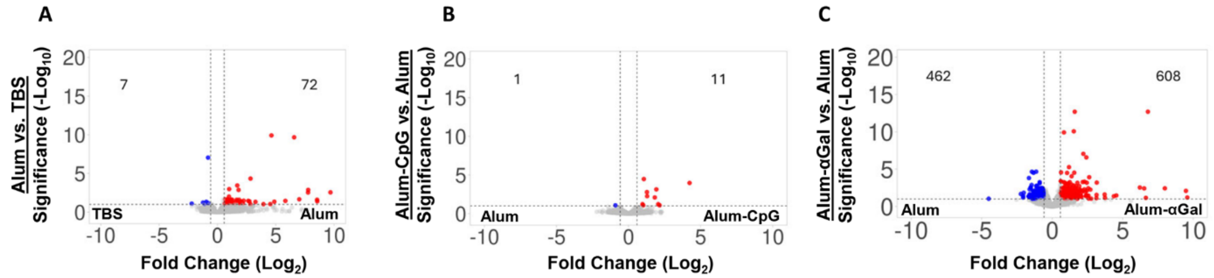

**Figure S1. The differential gene expressions in the mice inoculated with Alum, Alum-CpG, or Alum-αGal formulated CspZ-YA$_{C187S}$ vs. TBS.** Five pre-adolescent C3H/HeN mice per group were inoculated with TBS or CspZ-YA$_{C187S}$ formulated with Alum, Alum-CpG, and Alum-αGal at 0 and 14dpii in the fashion described in **Fig. 1A.** At 28dpii, the expression levels of genes in the spleen from each group of the mice were determined. Differentially expressed genes (DEGs) are defined by an adjusted P values less than 0.05 and absolute (log$_2$fold change) ≥0.58 or ≤0.58. The DEGs in the mice inoculated with CspZ-YA$_{C187S}$ formulated with **(A)** Alum vs. TBS, **(B)** Alum-CpG vs. TBS, or **(C)** Alum-αGal vs. TBS were plotted as a Volcano plots. The up- and down-regulated DEGs were shown in red and blue, respectively. The details of the immune-related DEGs were shown in **Dataset S1**.

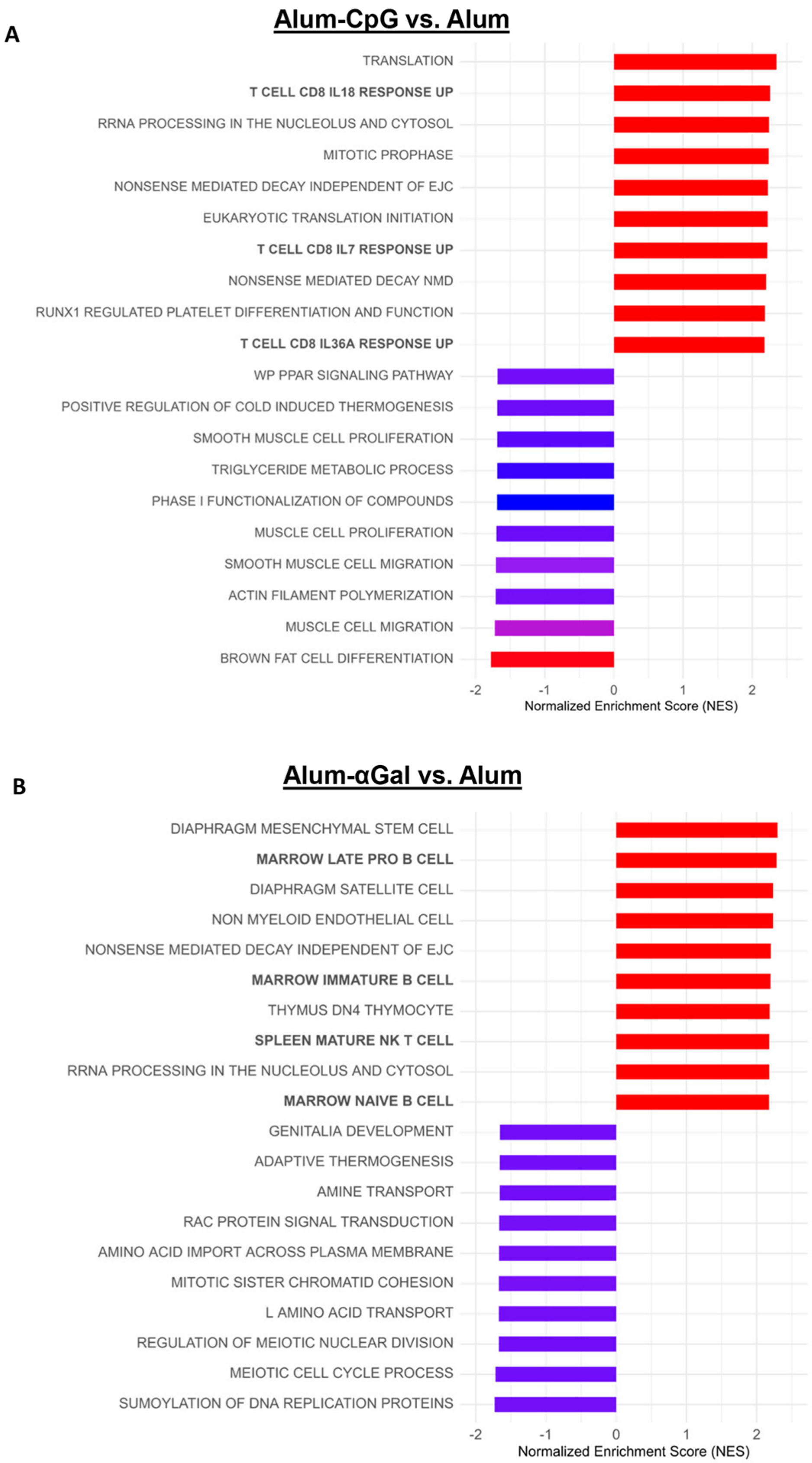
A
Alum-CpG vs. Alum
TRANSLATION
T CELL CD8 IL18 RESPONSE UP
RRNA PROCESSING IN THE NUCLEOLUS AND CYTOSOL
MITOTIC PROPHASE
NONSENSE MEDIATED DECAY INDEPENDENT OF EJC
EUKARYOTIC TRANSLATION INITIATION
T CELL CD8 IL7 RESPONSE UP
NONSENSE MEDIATED DECAY NMD
RUNX1 REGULATED PLATELET DIFFERENTIATION AND FUNCTION
T CELL CD8 IL36A RESPONSE UP
WP PPAR SIGNALING PATHWAY
POSITIVE REGULATION OF COLD INDUCED THERMOGENESIS
SMOOTH MUSCLE CELL PROLIFERATION
TRIGLYCERIDE METABOLIC PROCESS
PHASE I FUNCTIONALIZATION OF COMPOUNDS
MUSCLE CELL PROLIFERATION
SMOOTH MUSCLE CELL MIGRATION
ACTIN FILAMENT POLYMERIZATION
MUSCLE CELL MIGRATION
BROWN FAT CELL DIFFERENTIATION
-2
-1
0
1
2
Normalized Enrichment Score (NES)
B
Alum-αGal vs. Alum
DIAPHRAGM MESENCHYMAL STEM CELL
MARROW LATE PRO B CELL
DIAPHRAGM SATELLITE CELL
NON MYELOID ENDOTHELIAL CELL
NONSENSE MEDIATED DECAY INDEPENDENT OF EJC
MARROW IMMATURE B CELL
THYMUS DN4 THYMOCYTE
SPLEEN MATURE NK T CELL
RRNA PROCESSING IN THE NUCLEOLUS AND CYTOSOL
MARROW NAIVE B CELL
GENITALIA DEVELOPMENT
ADAPTIVE THERMOGENESIS
AMINE TRANSPORT
RAC PROTEIN SIGNAL TRANSDUCTION
AMINO ACID IMPORT ACROSS PLASMA MEMBRANE
MITOTIC SISTER CHROMATID COHESION
L AMINO ACID TRANSPORT
REGULATION OF MEIOTIC NUCLEAR DIVISION
MEIOTIC CELL CYCLE PROCESS
SUMOYLATION OF DNA REPLICATION PROTEINS
-2
-1
0
1
2
Normalized Enrichment Score (NES)

**Figure S2. GSEA analysis of the regulated genes in comparison of vaccination with CspZ-YA$_{C187S}$ formulated with Alum-CpG vs. Alum or Alum-αGal vs. Alum.** The top 10 enriched and pathways regulated in comparison of spleens from the mice immunized twice (0 and 14dpii) with CspZ-YAC187S formulated with Alum-CpG vs. Alum or Alum αGal vs. Alum are shown. Normalized enrichment scores (NES) and FDR q-value were determined by the GSEA software, color coeded and indicated within each enrichment plot.

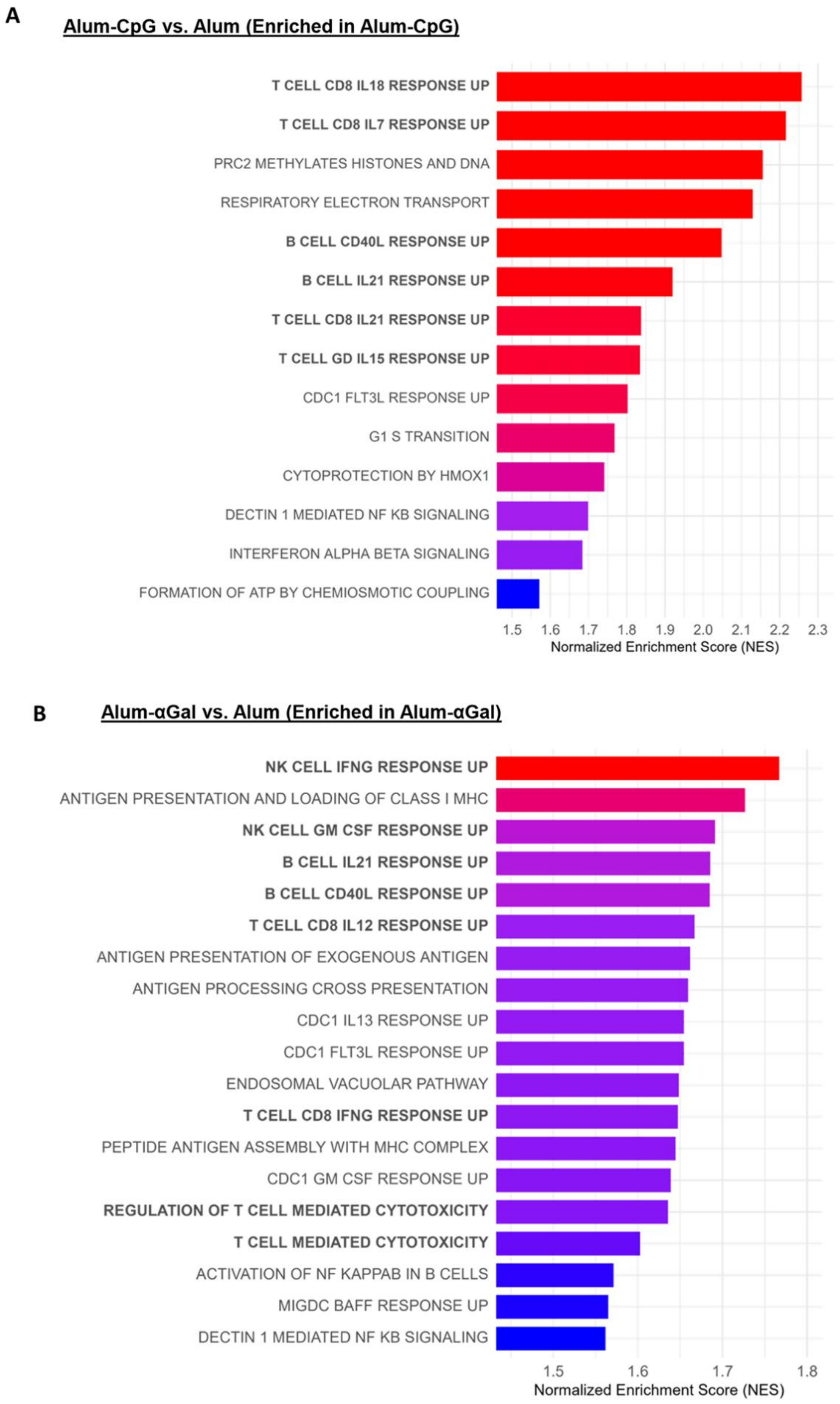
A
Alum-CpG vs. Alum (Enriched in Alum-CpG)
T CELL CD8 IL18 RESPONSE UP
T CELL CD8 IL7 RESPONSE UP
PRC2 METHYLATES HISTONES AND DNA
RESPIRATORY ELECTRON TRANSPORT
B CELL CD40L RESPONSE UP
B CELL IL21 RESPONSE UP
T CELL CD8 IL21 RESPONSE UP
T CELL GD IL15 RESPONSE UP
CDC1 FLT3L RESPONSE UP
G1 S TRANSITION
CYTOPROTECTION BY HMOX1
DECTIN 1 MEDIATED NF KB SIGNALING
INTERFERON ALPHA BETA SIGNALING
FORMATION OF ATP BY CHEMIOSMOTIC COUPLING
1.5 1.6 1.7 1.8 1.9 2.0 2.1 2.2 2.3
Normalized Enrichment Score (NES)
B
Alum-αGal vs. Alum (Enriched in Alum-αGal)
NK CELL IFNG RESPONSE UP
ANTIGEN PRESENTATION AND LOADING OF CLASS I MHC
NK CELL GM CSF RESPONSE UP
B CELL IL21 RESPONSE UP
B CELL CD40L RESPONSE UP
T CELL CD8 IL12 RESPONSE UP
ANTIGEN PRESENTATION OF EXOGENOUS ANTIGEN
ANTIGEN PROCESSING CROSS PRESENTATION
CDC1 IL13 RESPONSE UP
CDC1 FLT3L RESPONSE UP
ENDOSOMAL VACUOLAR PATHWAY
T CELL CD8 IFNG RESPONSE UP
PEPTIDE ANTIGEN ASSEMBLY WITH MHC COMPLEX
CDC1 GM CSF RESPONSE UP
REGULATION OF T CELL MEDIATED CYTOTOXICITY
T CELL MEDIATED CYTOTOXICITY
ACTIVATION OF NF KAPPAB IN B CELLS
MIGDC BAFF RESPONSE UP
DECTIN 1 MEDIATED NF KB SIGNALING
1.5 1.6 1.7 1.8
Normalized Enrichment Score (NES)

**Figure S3. GSEA analysis of the regulated genes in comparison of vaccination with CspZ-YA$_{C187S}$ formulated with Alum-CpG vs. Alum or Alum-αGal vs. Alum.** Five pre-adolescent C3H/HeN mice per group were inoculated with TBS or CspZ-YA$_{C187S}$ formulated with Alum, Alum-CpG, and Alum-αGal twice in the fashion described in **Fig. 1A.** GSEA analysis to identify the enriched pathways under the treatment of **(A)** Alum-CpG formulated or **(B)** Alum-αGal formulated CspZ-YA$_{C187S}$ in the comparison of Alum treated groups of mice.

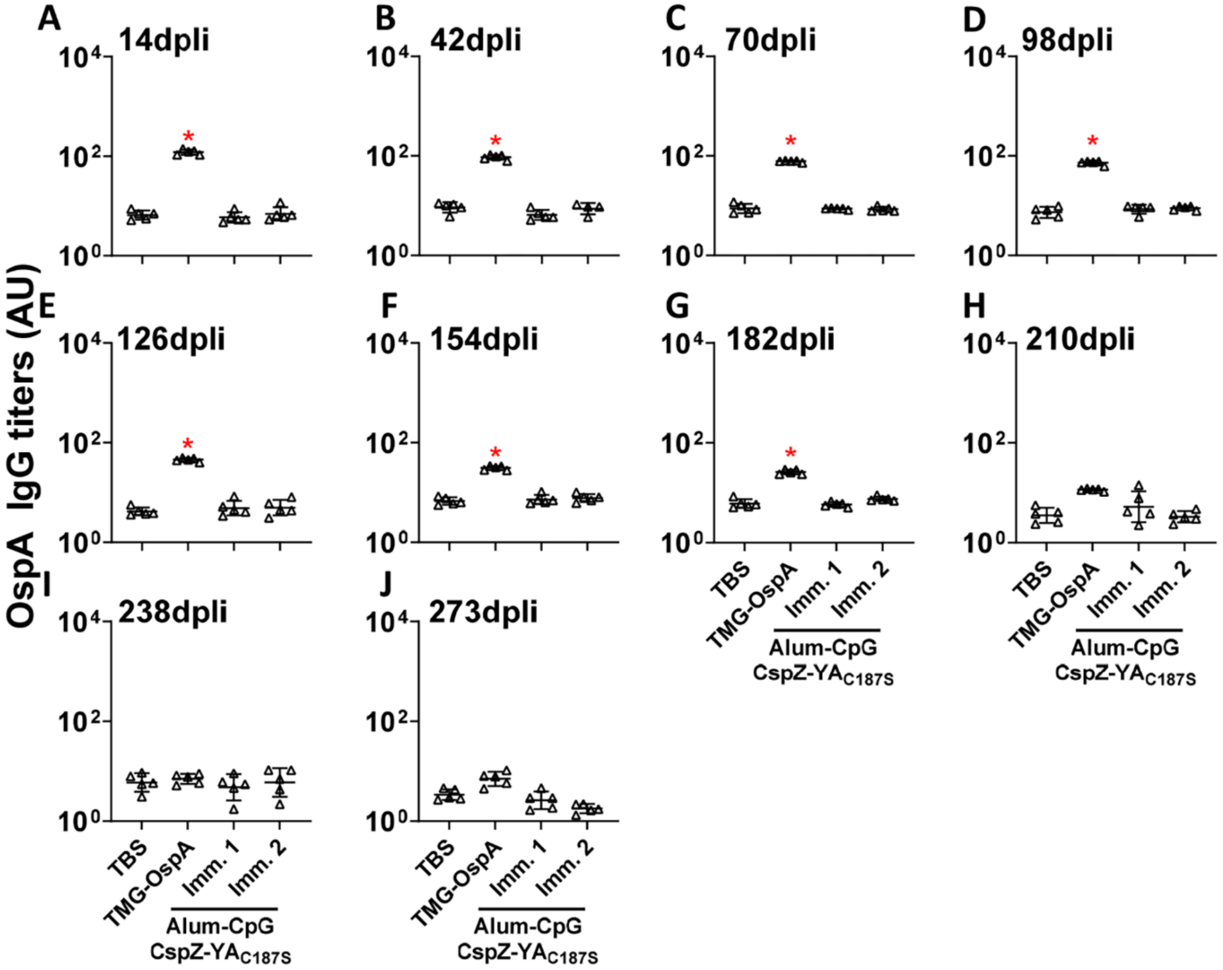

**Figure S4. The levels of anti-OspA IgG titers determined in the mice immunized with OspA or CspZ-YA$_{C187S}$ formulated with different adjuvants from 14 to 273 dpli.** Five pre-adolescent C3H/HeN receiving at 0 (Imm. 1) or both 0 and 14 (Imm. 2) dpii with CspZ-YA$_{C187S}$ formulated with Alum-CpG. Mice inoculated with TBS at 0 and 14 dpii, or TMG-formulated OspA at 0, 14, and 28dpii were included as control. Sera were collected at **(A)** 14, **(B)** 42, **(C)** 70, **(D)** 98, **(E)** 126, **(F)** 154, **(G)** 182, **(H)** 210, **(I)** 238, and **(J)** 273 dpli. The levels of total IgG against OspA in the sera were determined using quantitative ELISA. Data shown are the geometric mean ± geometric standard deviation of the titers of anti-OspA IgG from five mice per group. Statistical significance ($p < 0.05$, Kruskal Wallis test with the two-stage step-up method of Benjamini, Krieger, and

Yekutieli) of greater anti-OspA IgG titers in the indicated group were shown to be compared with the titers from TBS-inoculated mice (“*”).

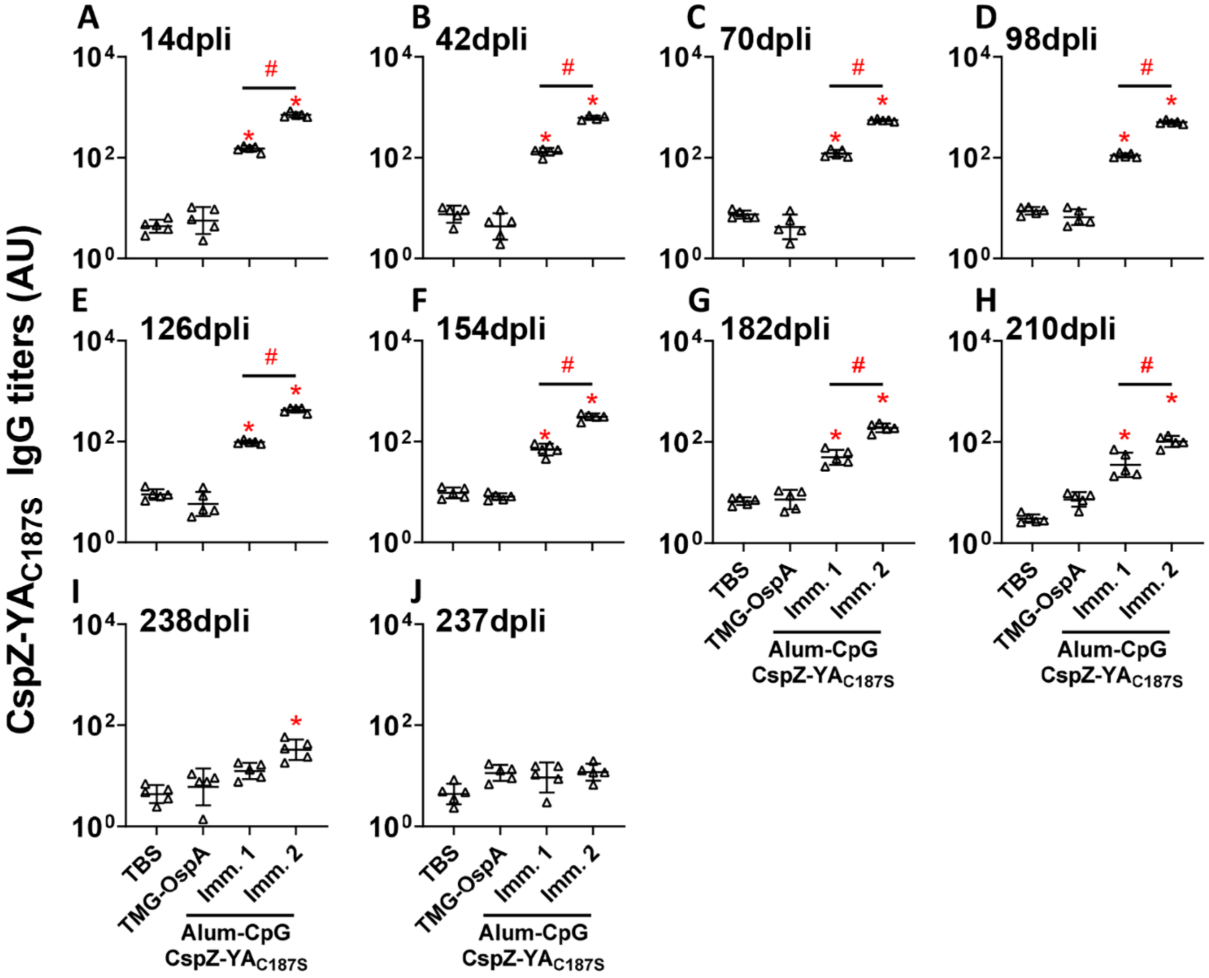

**Figure S5. The levels of anti-CspZ-YA$_{C187S}$ IgG titers determined in the mice immunized with OspA or CspZ-YA$_{C187S}$ formulated with different adjuvants from 14 to 273 dpli.** Five pre-adolescent C3H/HeN receiving at 0 (Imm. 1) or both 0 and 14 (Imm. 2) dpii with CspZ-YA$_{C187S}$ formulated with Alum-CpG. Mice inoculated with TBS at 0 and 14 dpii, or TMG-formulated OspA at 0, 14, and 28dpii were included as control. Sera were collected at **(A)** 14, **(B)** 42, **(C)** 70, **(D)** 98, **(E)** 126, **(F)** 154, **(G)** 182, **(H)** 210, **(I)** 238, and **(J)** 273 dpli. The levels of total IgG against CspZ-YA$_{C187S}$ in the sera were determined using quantitative ELISA. Data shown are the geometric mean ± geometric standard deviation of the titers of anti-CspZ-YA$_{C187S}$ antibodies from five mice per group. Statistical significance ($p < 0.05$, Kruskal Wallis test with the two-stage step-up method of Benjamini, Krieger, and Yekutieli) of greater anti-CspZ-$YA_{C187S}$ IgG in the indicated group were shown to be compared with the titers from TBS-inoculated mice ("*") or between indicated groups ("#").

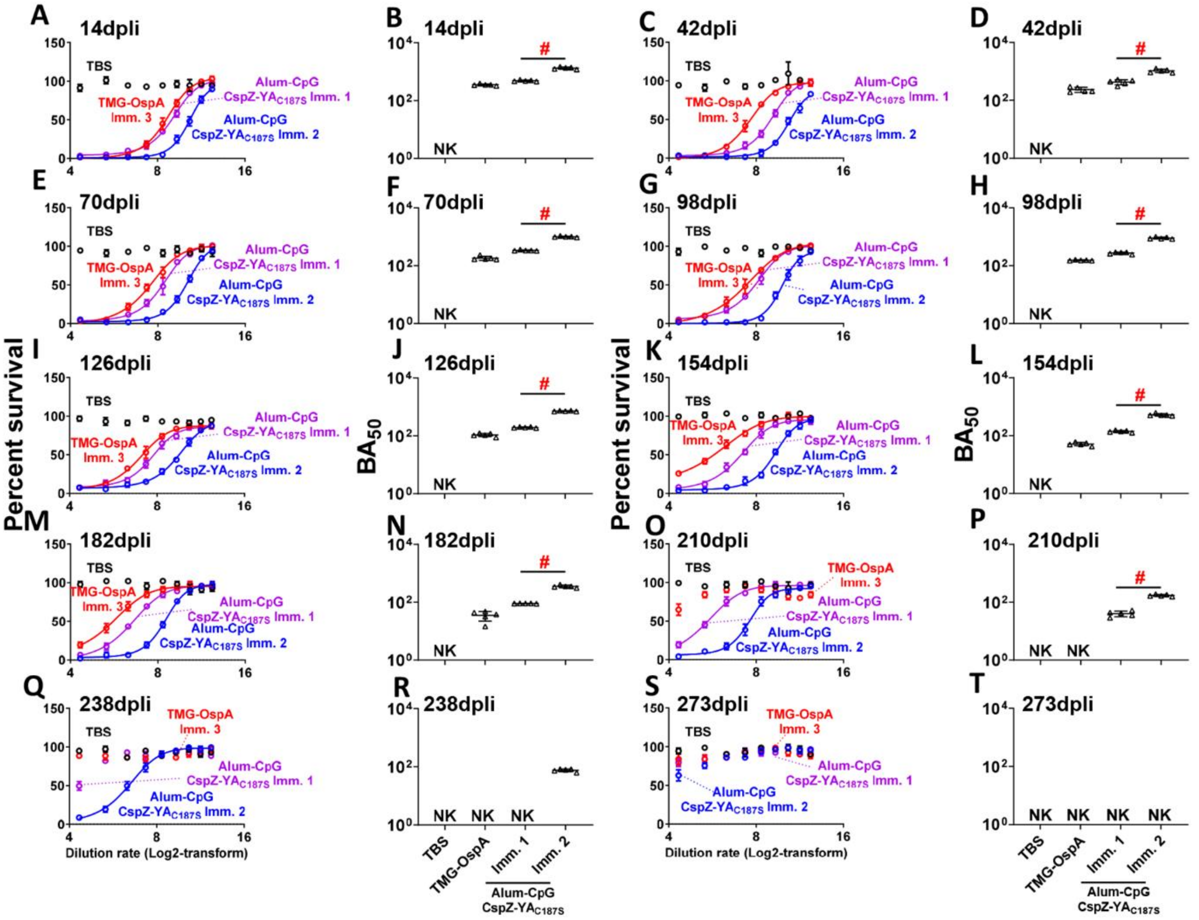

**Figure S6. The levels of bactericidal antibodies determined in the mice immunized with OspA or CspZ-YA$_{C187S}$ formulated with different adjuvants from 14 to 273 dpli.** Five pre-adolescent C3H/HeN receiving at 0 (Imm. 1), or both 0 and 14 (Imm. 2) dpii with CspZ-YA$_{C187S}$ formulated with Alum-CpG. Mice inoculated with TBS at 0 and 14 dpii, or TMG-formulated OspA at 0, 14, and 28dpii were included as control. Sera were collected at **(A-B)** 14, **(C-D)** 42, **(E-F)** 70, **(G-H)** 98, **(I-J)** 126, **(K-L)** 154, **(M-N)** 182, **(O-P)** 210, **(Q-R)** 238, and **(S-T)** 273 dpli. The sera were diluted as indicated, and mixed with guinea pig complement and *B. burgdorferi* B31-A3 for 24 hours. Surviving spirochetes were quantified from three fields of view microscopically in three independent experiments. **(A, C, E, G, I, K, M, O, Q, and S)** The survival percentage was derived from the proportion of serum-treated to untreated spirochetes. Data shown are the mean ± SEM of the survival percentage from three replicates in one representative experiment. **(B, D, F, H, J, L, N, P, R, and T)** The $BA_{50}$ value, representing the dilution rate that effectively killed 50% of spirochetes, was obtained from curve-fitting and extrapolation of Panel A, C, E, G, I, K, M, O, Q, and S for indicated time points. Data shown are the geometric mean ± geometric standard deviation of the borreliacidal titers from five mice per group in three experiments per mouse sample and shown in **Table S4.** Statistical significance ($p < 0.05$, Kruskal Wallis test with the two-stage step-up method of Benjamini, Krieger, and Yekutieli) of greater $BA_{50}$ was shown between indicated groups ("#").

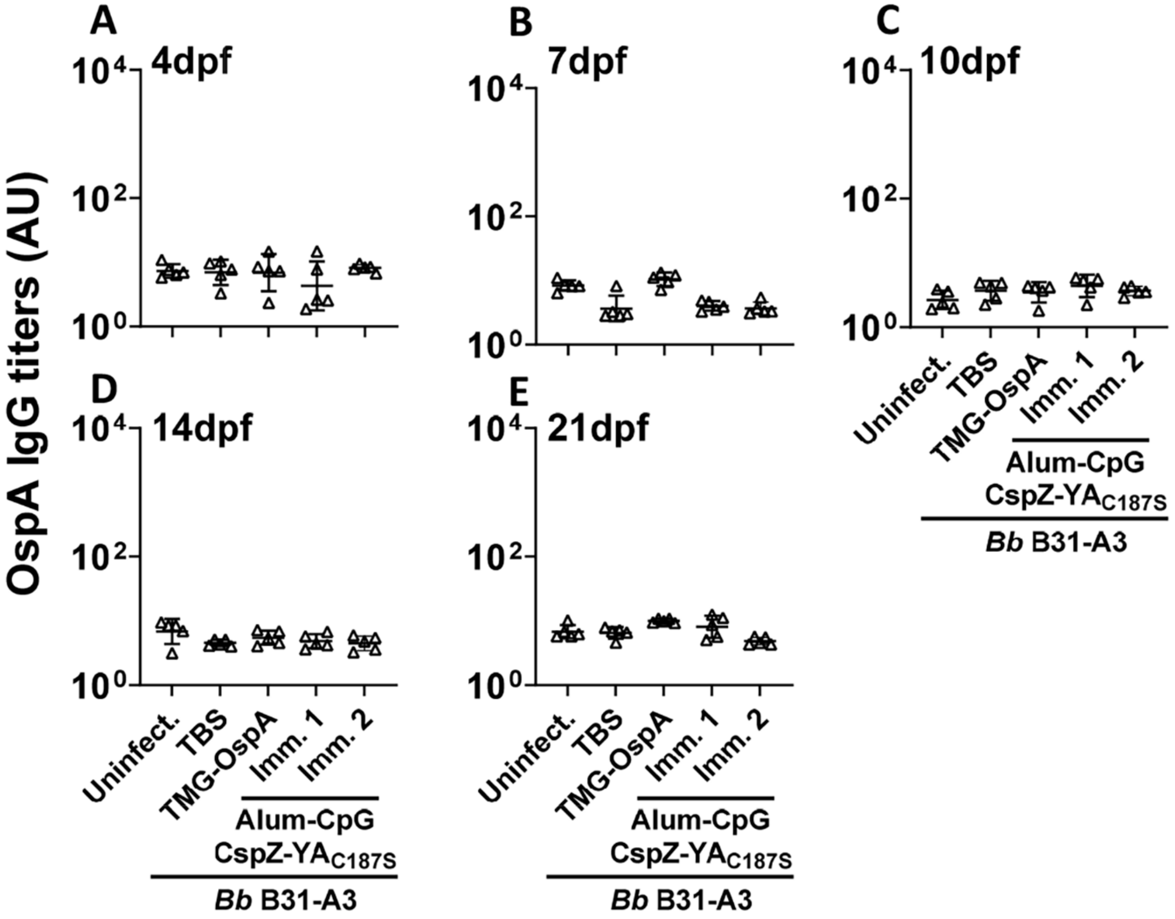

**Figure S7. The levels of anti-OspA IgG titers determined in the mice immunized with OspA or CspZ-YA$_{C187S}$ formulated with different adjuvants from 4 to 21 dpf.** Five pre-adolescent C3H/HeN receiving at 0 (Imm. 1), and both 0 and 14 (Imm. 2) dpii with CspZ-YA$_{C187S}$ formulated with Alum-CpG. Mice inoculated with TBS at 0 and 14 dpii, or TMG-formulated OspA at 0, 14, and 28dpii were included as control. The mice were infected using *I. scapularis* nymphs carrying *B. burgdorferi* strain B31-A3 at 273 dpli. Mice inoculated with TBS that are not fed on by nymphs were included as an uninfected control group (uninfect.). The sera were collected at **(A)** 4, **(B)** 7, **(C)** 10, **(D)** 14, and **(E)** 21 dpf. The levels of total IgG against OspA in the sera were determined using quantitative ELISA. Data shown are the geometric mean ± geometric standard deviation of the titers of anti-OspA IgG from five mice per group.

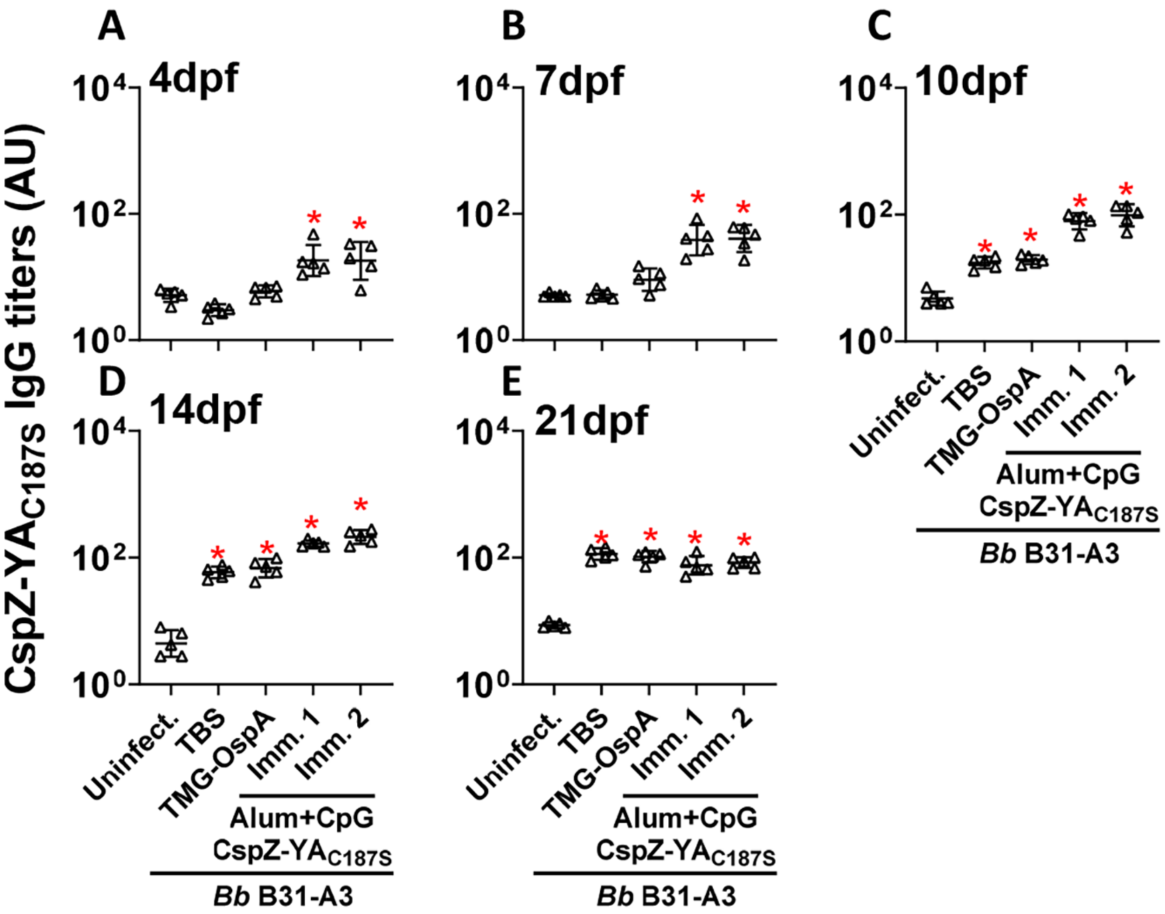

**Figure S8. The levels of anti-CspZ-YA$_{C187S}$ IgG titers determined in the mice immunized with OspA or CspZ-YA$_{C187S}$ formulated with different adjuvants from 4 to 21 dpf.** Five pre-adolescent C3H/HeN receiving at 0 (Imm. 1), and both 0 and 14 (Imm. 2) dpii with CspZ-YA$_{C187S}$ formulated with Alum-CpG. Mice inoculated with TBS at 0 and 14 dpii, or TMG-formulated OspA at 0, 14, and 28dpii were included as control. The mice were infected using *I. scapularis* nymphs carrying *B. burgdorferi* strain B31-A3 at 273 dpli. Mice inoculated with TBS that are not fed on by nymphs were included as an uninfected control group (uninfect.). The sera were collected at **(A)** 4, **(B)** 7, **(C)** 10, **(D)** 14, and **(E)** 21 dpf. The levels of total IgG against CspZ-YA$_{C187S}$ in the sera were determined using quantitative ELISA. Data shown are the geometric mean ± geometric standard deviation of the titers of anti-CspZ-$YA_{C187S}$ IgG from five mice per group. Statistical significance ($p < 0.05$, Kruskal Wallis test with the two-stage step-up method of Benjamini, Krieger, and Yekutieli) of greater anti-CspZ-$YA_{C187S}$ IgG titers in the indicated group were shown to be compared with the titers from uninfected mice ("*").

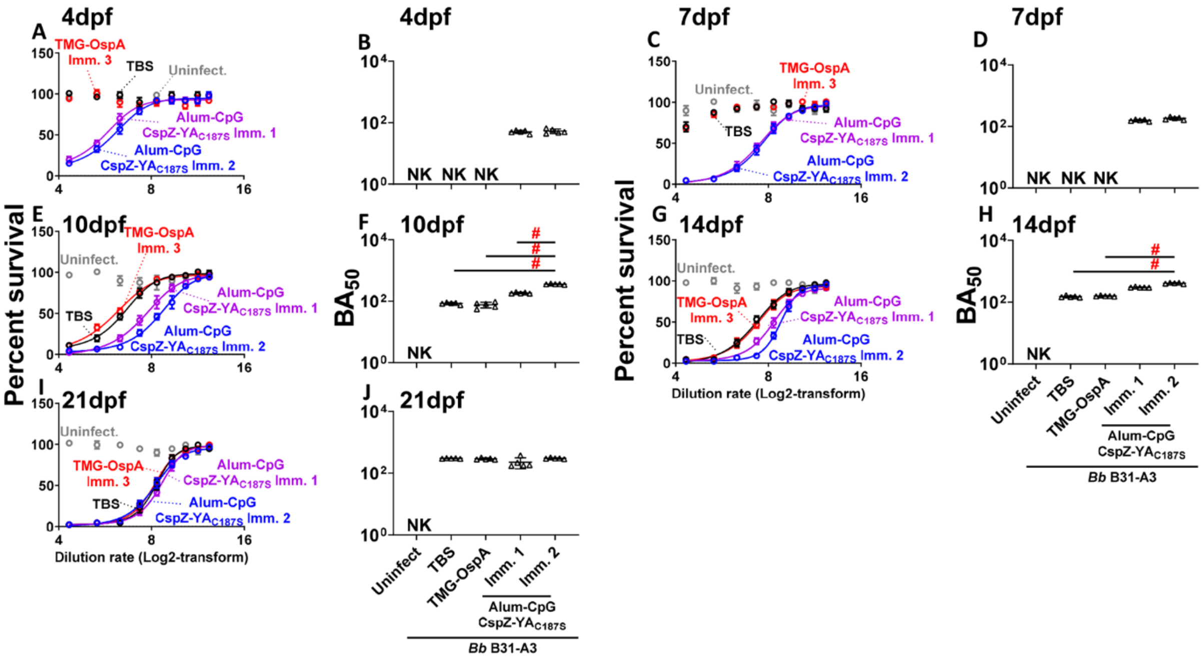

**Figure S9. The levels of bactericidal antibodies determined in the mice immunized with OspA or CspZ-YA$_{C187S}$ formulated with different adjuvants from 4 to 21 dpf.** Five pre-adolescent C3H/HeN receiving at 0 (Imm. 1), and both 0 and 14 (Imm. 2) dpii with CspZ-YA$_{C187S}$ formulated with Alum-CpG. Mice inoculated with TBS at 0 and 14 dpii, or TMG-formulated OspA at 0, 14, and 28dpii were included as control. The mice were infected using *I. scapularis* nymphs carrying *B. burgdorferi* strain B31-A3 at 273 dpli. Mice inoculated with TBS that are not fed on by nymphs were included as an uninfected control group (uninfect.). Sera were collected at **(A-B)** 4, **(C-D)** 7, **(E-F)** 10, **(G-H)** 14, and **(I-J)** 21 dpf. Sera were diluted as indicated, and mixed with guinea pig complement and *B. burgdorferi* B31-A3 for 24 hours. Surviving spirochetes were quantified from three fields of view microscopically in three independent experiments. **(A, C, E, G, and I)** The survival percentage was derived from the proportion of serum-treated to untreated spirochetes. Data shown are the mean ± SEM of the survival percentage from three replicates in one representative experiment. **(B, D, F, H, and J)** The BA$_{50}$ value, representing the dilution rate that effectively killed 50% of spirochetes, was obtained from curve-fitting and extrapolation of

Panel A, C, E, G, and I for indicated time points. Data shown are the geometric mean ± geometric standard deviation of the borreliacidal titers from five mice per group in three experiments per mouse sample and shown in **Table S5.** Statistical significance ($p < 0.05$, Kruskal Wallis test with the two-stage step-up method of Benjamini, Krieger, and Yekutieli) of greater $BA_{50}$ was shown between indicated groups (“#”).

### SUPPLEMENTAL TABLES

**Table S1. Bactericidal activities ($BA_{50}$) of sera from mice immunized with CspZ-$YA_{C187S}$ formulated with different adjuvants at 14-days post last inoculation.**

| Immun. frequency | $BA_{50}$ | | | | | |
|---|---|---|---|---|---|---|
| | TBS | CspZ-$YA_{C187S}$ | | | | |
| | | TMG | PAM3CSK4 | Alum | Alum+CpG | Alum+αGal |
| **Inoc. Once**[a] | N.K.[c] | 29±5 | N.K. | N.K. | 336±45 | 345±35 |
| **Inoc. Twice**[b] | N.K. | 220±17 | 111±38 | 87±8 | 1541±159 | 1989±555 |

[a]Inoculated at 0-day post initial inoculation
[b]Inoculated at 0- and 14-days post initial inoculation
[c]N.K. no killing was detected.

**Table S2. DESeq2 results for immune-related DEGs from the spleen of mice inoculated with CspZ-YA$_{C187S}$ formulated with Alum-CpG vs. Alum at 14-days post initial inoculation.**

| Gene | Adjusted P-values | Log2-fold change | Level ID |
|---|---|---|---|
| **Upregulated** | | | |
| Ighv1-70 | 0.075715 | 1.473107 | Immunoglobulin heavy variable 1-70 |
| Ighv8-12 | 0.084884 | 0.884879 | Immunoglobulin heavy variable V8-12 |
| Gdf6 | 0.077917 | 0.583196 | Growth differentiation factor 6 |
| **Downregulated** | | | |
| Cd34 | 0.078593 | -0.68092 | CD34 antigen |
| Klf9 | 0.039125 | -0.70214 | Kruppel-like transcription factor 9 |
| Adamts1 | 0.031401 | -0.71453 | ADAM metallopeptidase with thrombospondin type 1 motif1 |
| Cav2 | 0.075715 | -0.71715 | Caveolin 2 |
| Clec10a | 0.013235 | -0.98815 | C-type lectin domain family 10, member A |
| Errfi1 | 0.004227 | -1.00432 | ERBB receptor feedback inhibitor 1 |
| Mmp3 | 0.075715 | -1.05465 | Matrix metallopeptidase 3 |
| Pde1a | 0.039619 | -1.11783 | Phosphodiesterase 1A, calmodulin-dependent |
| Maob | 0.076815 | -1.47116 | Monoamine oxidase B |
| Alox15 | 0.015871 | -1.95182 | Arachidonate 15-lipoxygenase |
| Ebf3 | 0.077917 | -2.2296 | Early B cell factor 3 |
| Ccl11 | 0.076721 | -2.28078 | C-C motif chemokine ligand 11 |
| Serpina3c | 0.066586 | -2.71692 | Serine (or cysteine) peptidase inhibitor, clade A, member 3C |
| 9330159F19Rik | 0.033900 | -3.22287 | RIKEN cDNA 9330159F19 gene |

**Table S3. DESeq2 results for immune-related DEGs from the spleen of mice inoculated with CspZ-YA$_{C187S}$ formulated with Alum-αGal vs. Alum at 14-days post initial inoculation.**

| Gene | Adjusted P-values | Log2-fold change | Level ID |
|---|---|---|---|
| **Upregulated** | | | |
| Ighv8-12 | 0.092697 | 1.451218 | Immunoglobulin heavy variable V8-12 |
| H2-Q3 | 0.052531 | 1.413358 | Histocompatibility 2, Q region locus 3 |
| Igkv7-33 | 0.068092 | 1.119412 | Immunoglobulin kappa chain variable 7-33 |
| Socs1 | 0.056526 | 1.042779 | Suppressor of cytokine signaling 1 |
| Iglj3 | 0.085104 | 1.032291 | Immunoglobulin lambda joining 3 |
| Igkv12-46 | 0.068092 | 0.950486 | Immunoglobulin kappa variable 12-46 |
| H2-Q6 | 0.095829 | 0.935259 | Histocompatibility 2, Q region locus 6 |
| Ighv3-8 | 0.09962 | 0.860866 | Immunoglobulin heavy variable V3-8 |
| H2-Q2 | 0.008905 | 0.831371 | Histocompatibility 2, Q region locus 2 |
| Igkv8-19 | 0.068092 | 0.76336 | Immunoglobulin kappa variable 8-19 |
| Atoh8 | 0.022353 | 0.754916 | Atonal bHLH transcription factor 8 |
| Cd3d | 0.068092 | 0.67544 | CD3 antigen, delta polypeptide |
| Igkv2-109 | 0.052531 | 0.669317 | Immunoglobulin kappa variable 2-109 |
| H2-Q1 | 0.049911 | 0.668427 | Histocompatibility 2, Q region locus 1 |
| Iglc2 | 0.031594 | 0.642397 | Immunoglobulin lambda constant 2 |
| Tlr12 | 0.068092 | 0.639465 | Toll-like receptor 12 |
| **Downregulated** | | | |
| Siglec1 | 0.068092 | -0.61518 | Sialic acid binding Ig-like lectin 1, sialoadhesin |
| Ifit1 | 0.061232 | -0.62627 | Interferon-induced protein with tetratricopeptide repeats 1 |
| Rock 1 | 0.068709 | -0.64311 | Rho-associated coiled-coil containing protein kinase 1 |
| Pla2g4c | 0.088261 | -0.71386 | Phospholipidase A2, group IVC |
| Cpeb3 | 0.096371 | -0.74999 | Cytoplasmic polyadenylation element binding protein 3 |
| Cpeb4 | 0.095613 | -0.77284 | Cytoplasmic polyadenylation element binding protein 3 |

| | | | |
|---|---|---|---|
| Chp2 | 0.068092 | -0.84302 | Calcineurin-like EF hand protein 2 |
| Igkv11-125 | 0.043424 | -0.84604 | Immunoglobulin kappa variable 11-125 |
| Mmp27 | 0.092894 | -1.14547 | Matrix metallopeptidase 27 |

**Table S4. The bactericidal activities ($BA_{50}$) from the mice immunized with CspZ-$YA_{C187S}$ formulated with different adjuvants from 14 to 273-days post last inoculation.**

| Sample collecting time points (days post last inoc.) | $BA_{50}$ | | | |
|---|---|---|---|---|
| | TBS | OspA | CspZ-$YA_{C187S}$ | |
| | | TMG | Alum+CpG | |
| | | Inoc. Three times[a] | Inoc. Once[b] | Inoc. Twice[c] |
| **14** | N.K.[d] | 352±24 | 486±26 | 1328±109 |
| **42** | N.K. | 235±46 | 429±84 | 1042±114 |
| **70** | N.K. | 182±28 | 338±13 | 1003±104 |
| **98** | N.K. | 154±5 | 273±21 | 904±10 |
| **126** | N.K. | 107±14 | 195±7 | 719±10 |
| **154** | N.K. | 51±6 | 139±11 | 520±11 |
| **182** | N.K. | 35±13 | 90±12 | 351±11 |
| **210** | N.K. | N.K. | 40±8 | 171±10 |
| **238** | N.K. | N.K. | N.K. | 75±1 |
| **273** | N.K. | N.K. | N.K. | N.K. |

[a]Inoculated at 0-, 14-, and 28-days post initial inoculation
[b]Inoculated at 0-day post initial inoculation
[c]Inoculated at 0- and 14-days post initial inoculation
[c]N.K. no killing was detected.

**Table S5. The bactericidal activities ($BA_{50}$) from the mice immunized with CspZ-$YA_{C187S}$ formulated with different adjuvants from 0 to 21-days post nymph feeding.**

| Sample collecting time points (days after nymph feeding) | $BA_{50}$ | | | | |
|---|---|---|---|---|---|
| | Uninfect. | TBS | OspA | CspZ-$YA_{C187S}$ | |
| | | | TMG | Alum+CpG | |
| | | | Inoc. Three times[a] | Inoc. Once[b] | Inoc. Twice[c] |
| **4** | N.K.[d] | N.K. | N.K. | 50±5 | 52±1 |
| **7** | N.K. | N.K. | N.K. | 157±11 | 182±11 |
| **10** | N.K. | 85±6 | 76±16 | 185±8 | 358±10 |
| **14** | N.K. | 151±10 | 157±5 | 303±9 | 411±10 |
| **21** | N.K. | 300±7 | 291±19 | 225±87 | 301±10 |

[a]Inoculated at 0-, 14-, and 28-days post initial inoculation
[b]Inoculated at 0-day post initial inoculation
cInoculated at 0- and 14-days post initial inoculation
[c]N.K. no killing was detected.

**Table S6. *B. burgdorferi* and *E. coli* Strains used in this study.**

| Strain or plasmid | Genotype or characteristic | Source |
|---|---|---|
| <u>*B. burgdorferi*</u> | | |
| B31-A3 | Clone A3 of *B. burgdorferi* B31 isolated from *I. scapularis* ticks in US. | [1] |
| <u>*E. coli*</u> | | |
| BL21(DE3)/pET41a-CspZ-YA$_{C187S}$ | BL21(DE3) producing residues 19 to 237 of CspZ with tyrosine-207 and tyrosine-211 replaced by alanine, and cysteine-187 replaced by serine | [2] |

### SUPPLEMENTAL DATASET

**Dataset S1. All differentially expressed genes from the spleen of mice inoculated with CspZ-YA$_{C187S}$ formulated with different adjuvants.** At 14-days post last immunization, shown are the differential expressed genes and their functions in the mice inoculated with CspZ-YA$_{C187S}$ formulated with (**Sheet 1**) Alum vs. TBS, (**Sheet 2**) Alum-CpG vs. TBS, (**Sheet 3**) Alum-αGal vs. TBS, (**Sheet 4**) Alum-CpG vs. Alum, and (**Sheet 5**) Alum-αGal vs. Alum.
